# PI(3)P signaling regulates endosomal flux underlying developmental synaptic remodeling via Rab4

**DOI:** 10.1101/2025.02.07.636819

**Authors:** Kamaldeep Singh, Semanti Das, Dipti Rai, Sabyasachi Sutradhar, Asmita Sarkar, Jonathon Howard, Krishanu Ray

**Author notes:** Cell Biology, Department of Biology, Faculty of Science, Utrecht University, Utrecht, the Netherlands (Semanti Das, Dipti Rai).

## Abstract

Rab4 GTPase, essential for endosomal sorting and trafficking, is implicated in synaptic atrophy and dementia. To uncover the underlying mechanism, we studied the correlation between Rab4 vesicle transport in axons and episodic remodeling of synapses in the central nervous system (CNS) of *Drosophila* larvae. We found that synapse-bound traffic and presynaptic enrichment of Rab4 vesicles increase during the programmed, transient contraction of synapses in the ventral neuropil region at a specific larval stage. This coincides with the episodic activation of insulin and Vps34-mediated signaling, which elevates phosphatidylinositol-3-phosphate levels on Rab4 vesicles. The presence of this phospholipid on Rab4-associated vesicles recruit a PX-domain-containing motor protein, Klp98A, accelerating synapse-directed traffic. This, in turn, increases presynaptic enrichment of Rab4 during the developmentally programmed synapse contraction phase. Our findings elucidate the molecular mechanism that regulates developmental synaptic plasticity in the CNS via insulin signaling and directed axonal transport of endosomes.

**Author summary:** Healthy brain function relies on the growth and remodeling of synaptic connections between nerve cells. This process involves carefully orchestrated movements of proteins and membranes inside the neurites. For instance, endosome movement in the nerve cell extensions helps guide neuronal growth, recycle parts of the synapse, and alter signal transmissions between neurons. Rab GTPases, a group of small G-proteins, are crucial for these processes. Here, we aimed to understand what triggers these processes and how the ’supply chain’ within nerve cell extensions is connected to changes at the synapse. In fruit fly larvae, we observed that the movement of Rab4 vesicles in the nerve cell extensions is inversely related to gross synapse content. A further examination revealed that onset of insulin signaling accelerates the Rab4 vesicles towards synapse through a lipid kinase called Vps34, which makes a signaling lipid called phospatidylinnositol-3-phosphate. Higher levels this lipid molecule on Rab4 vesicles attract a motor protein, Klp98A, speeding up their movement during the synaptic contraction phase. Our findings reveal a new mechanism that regulates Rab4 vesicle transport and its impact on synapse stability. The study further suggests potential targets for controlling age-related dementia likely to be caused by excessive insulin signaling in neurons.

## Introduction

Rab4 is a critical organizer of endosomal sorting and plays a crucial role in several processes like recycling of surface receptors [1], axon outgrowth [2], maintenance of spine morphology, and synaptic plasticity [3]. Rab4 activation promotes its recruitment on the endosomal membrane and activates vesicle trafficking through kinesin and dynein motors. Rab4-associated vesicles transport various cargos such as tetraspanins [4], neuregulins [5], integrins [6], and astrotactins [7], and regulates neuronal processes like trafficking and degradation of surface proteins, synapse formation, maturation, and maintenance. Unsurprisingly, mutations in some of these cargos are also implicated in neurodevelopmental [7] and psychiatric disorders [5]. Constitutive activation of Rab4 and its enrichment at the presynaptic terminals induces synaptic atrophy in *Drosophila* [8]. Further, brain autopsies of Alzheimer’s patients indicate an inverse correlation between the Rab4 levels in the cholinergic basal forebrain (CBF) and CA1 neurons of the hippocampus and their cognitive abilities [9,10]. Together, these observations suggest that the Rab4 activation in neurons must be tightly controlled to maintain synapse homeostasis. Therefore, understanding the yet unidentified molecular mechanisms regulating the Rab4 activation and the movement of Rab4-associated vesicles (henceforth called Rab4 vesicles) in neurons is essential for better comprehension of membrane trafficking functions in health and disease.

Besides the widely studied roles of insulin signaling in regulating metabolism [11], growth [12], and development [13], aberrant insulin signaling in the brain is also associated with cognitive decline with aging in *Drosophila* and humans [14,15]. Insulin signaling in the central nervous system (CNS) is implicated in amyloid precursor protein (APP) and huntingtin-associated protein 1 (HAP1) transport [16] and regulates AMPA receptor endocytosis and synaptic plasticity in hippocampal neurons [17]. However, the underlying mechanisms remain unclear. In addition, a few studies have also shown that insulin regulates intracellular transport by modulating the recruitment of microtubule-based motor proteins in peripheral tissues [18,19]. Specifically, insulin stimulation activates Rab4 and promotes its interaction with kinesin-2 to regulate GLUT4 recycling in adipocytes [18]. Therefore, we hypothesized that insulin signaling might also control synaptic stability by regulating the axonal transport of Rab4 vesicles.

To test this hypothesis, we first established a developmental system of periodic synapse remodeling in the CNS of 3^rd^ instar *Drosophila* larvae. It revealed that programmed contraction of synaptic content in the ventral neuromere coincides with episodic increase of Rab4 movement towards synapse and Rab4 enrichment at the presynaptic compartment. *In vivo* time-lapse imaging coupled with genetic and pharmacological perturbations showed that Dilp2 and dInR-mediated insulin signaling could accelerate the anterograde axonal transport of a subset of Rab4 vesicles in *Drosophila* cholinergic neurons. Further, a combined RNAi and chemical inhibitor screen of all known phosphatidylinositol-3-kinases (PI3Ks) in *Drosophila*, indicated a specific involvement of the Class-III PI3K, Vps34, in the process. We demonstrated that the activation of Vps34 by insulin signaling leads to the production of Phosphatidylinositol-3-Phosphate [PI(3)P] on Rab4 vesicles in the axon. As a consequence, Klp98A, a kinesin-3 family motor, is recruited by PI3P on Rab4 vesicles, activating their anterograde movement. Thus, this study elucidated a molecular basis for the acceleration of anterograde axonal transport of Rab4 vesicles in neurons by insulin signaling, which is likely to play a critical role in maintaining of synaptic homeostasis in vivo. Altogether, our study a) established an *in vivo* model system to investigate developmental synaptic plasticity in the CNS, b) provides a molecular mechanism by which neuronal insulin signaling regulates directional endosomal transport in the axons, and c), elucidates functional consequences of altered endosomal traffic on synaptic homeostasis in the CNS.

## Results

### The central nervous system (CNS) of *Drosophila* undergoes extensive synaptic remodeling during the third-instar larva stage

The size of *Drosophila* third-instar larvae and their CNS undergo significant mass and volume expansion essential for metamorphosis [20–22], providing a suitable model system to investigate developmental synaptic plasticity and its underlying mechanism. The larval CNS (Fig. 1A) consists of two large optic lobes connected by a suboesophageal ganglion (SOG) and a bilaterally symmetric ventral nerve cord (VNC) composed of an easily distinguishable cortex region rich in cell bodies surrounding the synapse-bearing core called the neuropil (Fig. 1B). VNC neuropil is divided into twelve segments (3 thoracic and 9 abdominal) along the anteroposterior body axis, with each segment comprising two bilaterally symmetric hemisegments. Each hemisegment consists of neurite arborizations and synapses contributed by segmentally organized sensory, motor and interneurons. It also contains neurite extensions form the anterior and posterior segments.

**Fig. 1.**
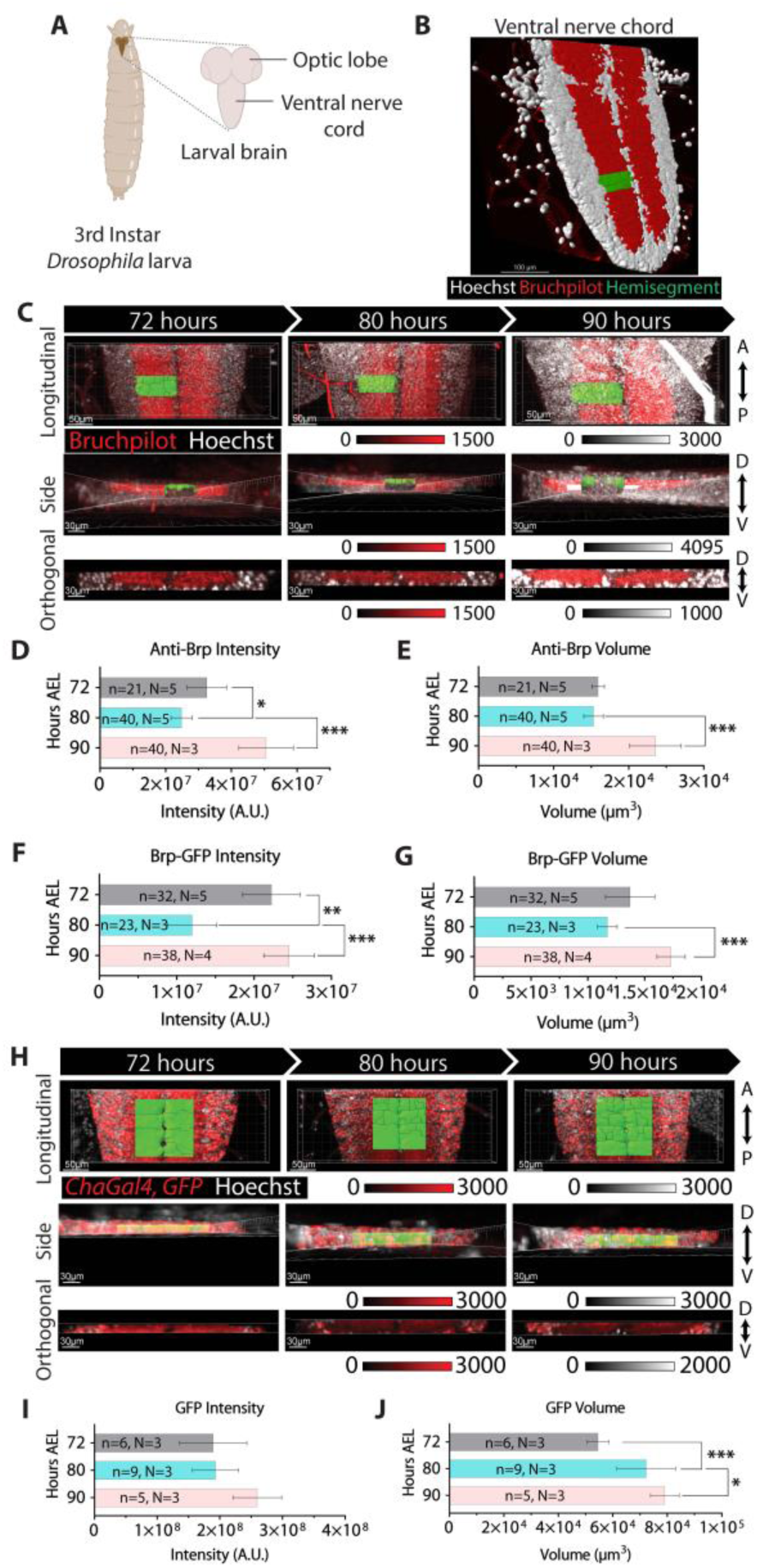
Synaptic remodeling in the ventral nerve cord during third-instar larval stage. **(A)** Schematic illustrates different parts of the 3rd instar larval brain. **(B)** 3D volume rendered image of a larval ventral nerve cord (VNC) stained with anti-Bruchpilot (green, presynaptic junctions) and Hoechst (red, cortex). Blue highlights a single neuromere hemisegment. **(C)** A3-A6 region (A5 highlighted in green is a ‘surface’ marked in Imaris®) of a larval VNC stained by anti-Bruchpilot (red) and Hoechst (white). **(D-G)** Mean ± SD of the volume of synaptic region and Bruchpilot enrichment in A3-A6 hemisegments marked by anti-Bruchpilot (D-E) and endogenous Bruchpilot-GFP (F-G) backgrounds. **(H)** A3-A6 segments (green box) marked by *chaGal4>GFP* (red) at 72-90 hours AEL. **(I-J)** Mean ± SD of GFP intensities and hemisegment volumes (I) and (J) of cholinergic neuromeres. The pairwise significance of the difference is estimated using one way ANOVA and Tukey’s correction for multiple comparisons.

Using 3D volume rendering of the confocal images of larval VNCs immunostained for Bruchpilot (Brp, an ELKS orthologue and a standard pre synaptic marker), we estimated synaptic volumes of A3-A6 hemisegments in progressively aged third-instar larvae. Here, A3-A6 hemisegments were selected for measurements due to negligible variation in synaptic volume and Bruchpilot enrichment amongst A3-A6 hemisegments at a given age (Fig. S1A-C). This analysis identified phases of periodic synaptic remodeling during 72-90 hours after egg laying (AEL). While the neuropil volume, marked by Brp/ELKS immunostaining and endogenously-expressed Brp-GFP, did not increase from 72-80 hours AEL, Brp enrichment decreased during this period (Fig. 1C-E and S1D). In contrast, both the synaptic volume and Brp enrichment increased from 80-90 hours AEL (Fig. 1C-E and S1D). Additionally, the total volume occupied by cholinergic neurites, estimated using tissue-specific expression of soluble GFP, increased from 72-90 hours AEL (Fig. 1F-H), indicating a progressive increase of neurite density and presynaptic volume during this entire developmental phase. Together, these data revealed that the neuropil expansion during 72-90 hours AEL is associated with programmed remodeling of synapses in the VNC.

### Presynaptic Rab4 enrichment and synaptic density loss in developing CNS correlate with increased anterograde axonal transport of Rab4 vesicles

Previous studies employing ectopic expression of constitutively active and dominant-negative Rab4 mutants revealed an inverse correlation between presynaptic Rab4 intensity and synapse volume in the third-instar larval CNS [8]. Estimations of presynaptic Rab4 density and Brp/ELKS density in the neuropil region between 72–90 hours AEL further confirmed an antiphase correlation between episodic changes in Rab4 accumulation at presynaptic sites and Brp/ELKS levels (Fig. 2A–B), suggesting that Rab4 enrichment at synapses may reduce synaptic contacts. Moreover, time-lapse imaging of cholinergic neurons expressing Rab4-mRFP at 72, 80, and 90 hours AEL (Fig 2C and movie S1) revealed a direct correlation between the anterograde flow of Rab4 vesicles and presynaptic Rab4 enrichment. The fraction of anterogradely moving Rab4 vesicles significantly increased from 72-80 hours AEL and then decresed from 80-90 hours AEL (Fig. 2D), seemingly at the expense of the retrograde fractions. This programmed increase in the anterograde fraction of Rab4 vesicles from 72-80 hours AEL could be promoted by: (a) enhanced anterograde velocity or processivity, (b) inhibition of retrograde movement, or (c) combination of both.

**Fig. 2.**
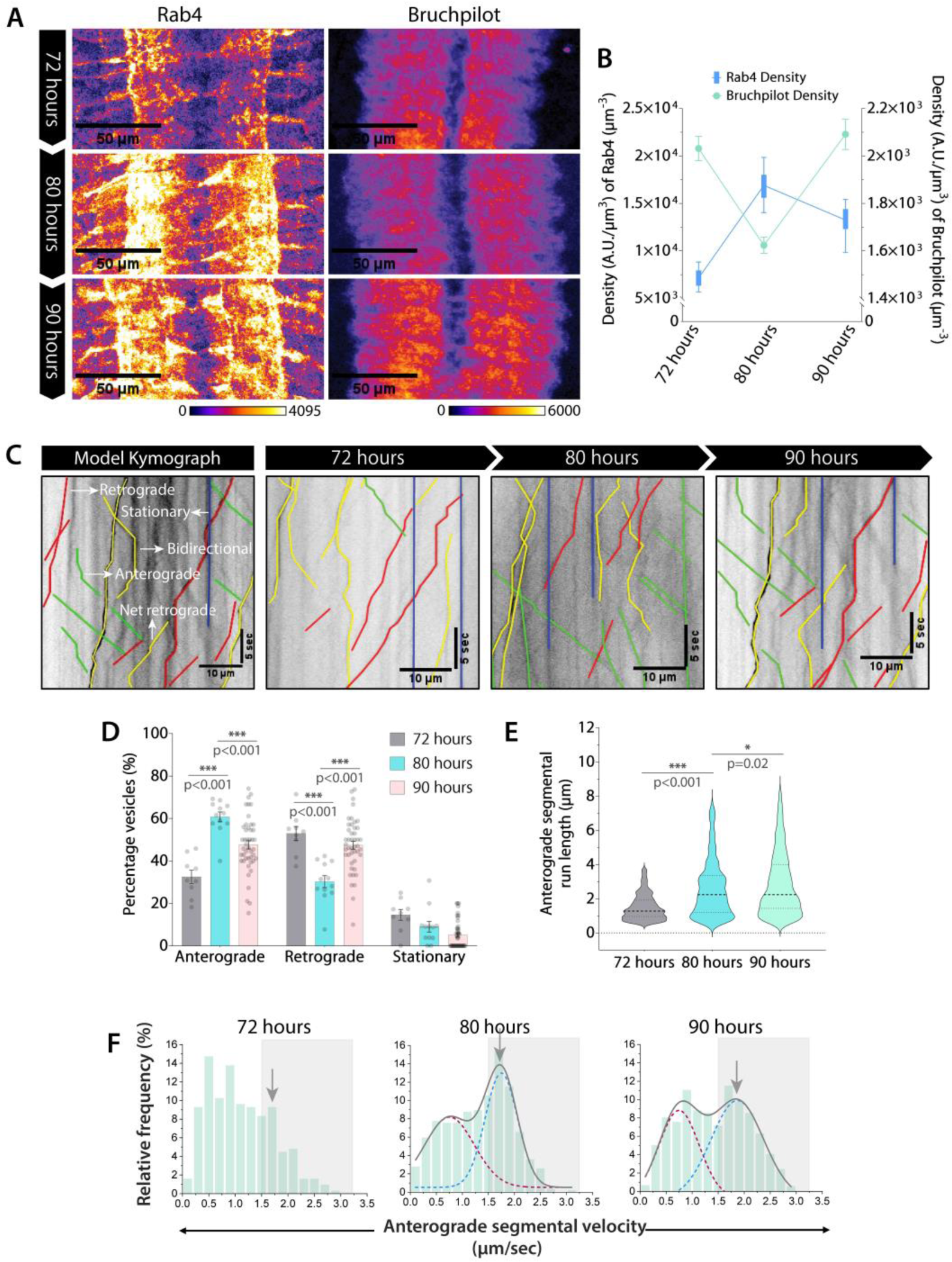
Synapse enrichment and axonal transport of Rab4 vesicles in developing larval VNC. **(A)** A3-A6 segments of VNC from 72-90 hours AEL stained with the Rab4 and Bruchpilot antibodies. Scale bar: 50 μm. **(B)** Rab4 (blue) and Bruchpilot (green) staining densities (A.U./μm^3^) in A3-A6 hemisegments (n>40, N=4-8 larvae). **(C)** Kymographs of Rab4 vesicles transport at 72-90 hours AEL. **(D-F)** Relative distribution (D, n≥9 segmental nerves, N = 3-5 larvae), anterograde segmental run length (E, n>300 runs), and anterograde segmental velocity (F) of Rab4 vesicles. The cumulative distribution (grey) as a sum of two Gaussians highlights slow (maroon) and fast-moving (blue) populations (See methods for details). Grey box marks the fast-moving (≥ 1.5 µm/sec) runs. The pairwise significance of difference was estimated using the Mann-Whitney U-test.

To investigate the underlying cause, we analyzed the transport parameters of individual segmental runs - specifically run length and velocity. The analysis revealed a significant increase in anterograde segmental run length of Rab4 vesicles from 72-80 hours AEL followed by a moderate increase from 80-90 hours AEL (Fig. 2E). Additionally, during 80 and 90 hours AEL, anterograde velocities of Rab4 vesicles exhibited a bimodal distribution, categorized into slow (0.0–1.5 µm/sec) and fast (1.5–3.0 µm/sec) moving segments (Fig. 2F). A similar bimodal distribution was also observed in the retrograde direction and was divided into slow (0.0 – 1.0 µm/sec) and fast-moving runs (1.0 – 2.25 µm/sec) (fig. S2B). The frequency of fast-moving anterograde runs increased significantly from 72-80 hours AEL followed by a significant reduction at 90 hours AEL (Fig. 2F and table S1; Kolmogorov Smirnov test, p<0.001). Finally, we observed a significant and progressive increase in the retrograde segmental run length and velocity of Rab4 vesicles from 72-90 hours AEL (fig. S2A, B and table S1; Kolmogorov Smirnov test, p<0.001).

Collectively, these results suggest that programmed alterations of the anterograde segmental velocity and run length during the developing third-instar larvae stage can change the anterograde fraction of Rab4 vesicles in the axons, which in turn may alter the Rab4 enrichment and reduce the synaptic density in the neuropil region. This observation raised a new question – what triggers the increase in the anterograde speed, run-length, and fraction of Rab4 vesicles during these phases?

### Cell-autonomous insulin signaling could selectively increase the anterograde fraction of Rab4 vesicles

Acute insulin stimulation activates Rab4 via type 1 PI3K, which recruits kinesin-2 on the recycling endosome marked by Rab4, thereby increasing the GLUT4 receptor levels in apical plasma membrane [18,23]. Rab4 activity, PI3K, and kinesin-2 motor are also implicated in Rab4 vesicle transport in axons [8]. Therefore, we hypothesized that insulin signaling could regulate Rab4 vesicle transport in axons. Insulin receptor (InR) is expressed in the sensory neurons [24] and is enriched in the axons of mechanosensory neurons of *Drosophila* pupa [25]. We further confirmed that InR-CFP ectopically expressed is cholinergic neurons is enriched both in the axons and the cortex region of VNC (fig. S3A-D).

To understand whether the InR-mediated signaling contributes to the axonal transport of Rab4 vesicles, we knocked down InR in the cholinergic neurons using two different UAS-RNAi lines known to suppress the expression of *Drosophila* insulin receptor (dInR). The InR knockdowns by both BL31594 (InR^RNAi^-1, Valium1, weak) and BL51518 (InR^RNAi^-2, Valium 20, strong) significantly reduced the anterograde fractions of Rab4 vesicles by ∼8% (InR^RNAi^-1) and ∼18% (InR^RNAi^-2), respectively, at 90 hours AEL (Fig. 3A). In comparison, overexpression of constitutively active insulin receptor (InR^CA^) significantly increased the anterograde fraction of Rab4 vesicles by ∼10% at 90 hours AEL (Fig. 3A), matching the level observed at 80 hours AEL (Fig. 2D). The increase in anterograde fraction upon overexpression of InR^CA^ was abolished by acute treatment with LY294002, which inhibits all known PI3Ks inside the cell (Fig. 3A). The data suggested that insulin signaling could act in a cell-autonomous manner, likely via PI3K, and the signaling dose could accordingly influence the anterograde movement of a subset of Rab4 vesicles in the axons.

**Fig. 3.**
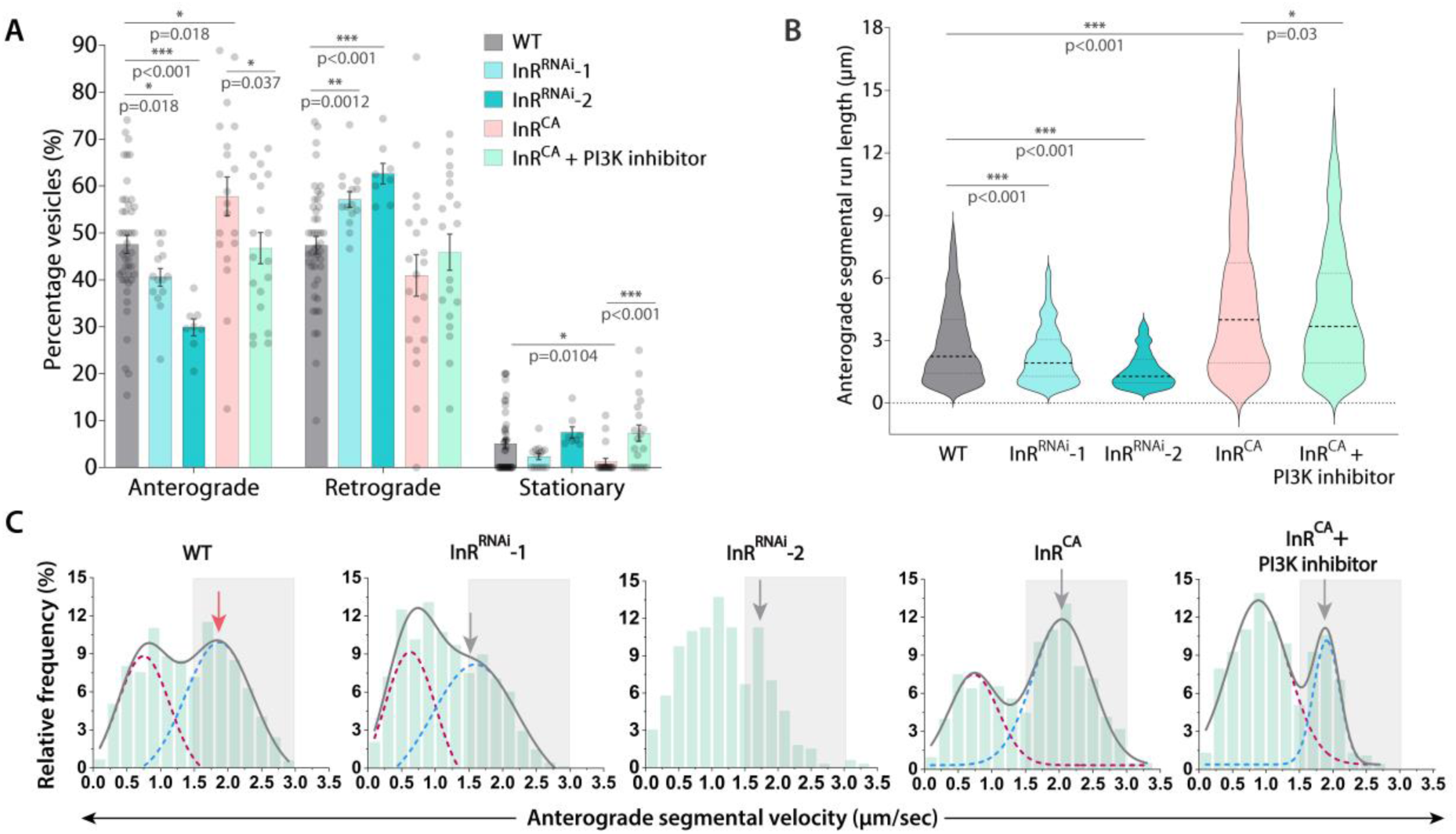
InR-signaling contributes to the axonal transport of Rab4 vesicles. **(A)** Relative distribution of the Rab4 vesicle movement (n≥9 segmental nerves, N = 3-5 larvae) in the wild-type control (WT), InR^RNAi^ and InR^CA^ overexpression backgrounds at 90 hours AEL. **(B, C)** Anterograde segmental run length (B) and segmental velocity distribution (C) of Rab4 vesicles in different genetic backgrounds (n>200 runs). The cumulative distribution (grey) as a sum of two Gaussians highlights slow (maroon) and fast-moving (blue) populations. Grey box marks the fast-moving (≥ 1.5 µm/sec) runs. The pairwise significance of difference was estimated using the Mann-Whitney U-test.

Further analysis of the motility data revealed a significant reduction in the anterograde segmental run length in both InR^RNAi^ backgrounds and increased in the InR^CA^ background (Fig. 3B). As expected, this InR activity-dependent increase in anterograde runs was partially suppressed by acute PI3K inhibition in the InR^CA^ background (Fig. 3B). A milder effect was observed on the retrograde run length of Rab4 vesicles (fig. S3E). Additionally, the frequency of fast-moving anterograde runs was reduced in both the InR^RNAi^-1/2 backgrounds and increased in the InR^CA^ overexpression background (∼15%; Fig. 3C and table S1). As expected, the LY294002 treatment in the InR^CA^ background suppressed the anterograde runs by ∼14% as compared to the untreated InR^CA^ preparations (Fig. 3C and table S1). Similar yet milder effects were observed on the frequency of fast-moving retrograde runs in the InR^RNAi^-1, InR^CA^, and LY294002-treated InR^CA^ backgrounds (fig. S3E and table S1).

Altogether, these observations suggest that the activation of cell-autonomous insulin signaling via dInR could increase the anterograde speed and processivity of Rab4 vesicles leading to a significant elevation of the anterograde flow of these vesicles in the cholinergic neurons of *Drosophila* (Fig. 2D). However, its effects on the retrograde movement of Rab4 vesicles were not conclusive.

### Treatment with human insulin and Dilp2 increases the anterograde fraction, velocity, and run-lengths of Rab4 vesicles

To understand the molecular mechanism by which insulin signaling activation could increase the anterograde velocity of Rab4 vesicles, we developed a pharmacological paradigm to stimulate the process in axons. Acute stimulation (15 minutes) with 1.7 nM human insulin in the bath (movie S3) significantly increased the proportion of anterogradely moving Rab4 vesicles by ∼10% and proportionately reduced the retrograde fraction (Fig. 4A). Of the eight *Drosophila* insulin-like peptides (Dilps) [26], Dilp2 and Dilp5 are the most abundantly expressed in the brain tissue [27], with Dilp2 having the highest homology (∼35% sequence identity) to human insulin [28]. Therefore, to understand the physiological relevance of the effect, we tested the effect of recombinant Dilp2 and Dilp5 on the axonal transport of Rab4 vesicles. Like human insulin, acute stimulation with 17 nM and 170nM Dilp2 also increased the net anterograde fraction of Rab4 vesicles (Fig. 4A). Whereas acute stimulation with Dilp5 even at 10-fold higher concentrations had no significant effect on the anterograde fraction of Rab4 vesicles, indicating a specificity in Dilp2 action (fig. S4A and table S1). These observations suggest that both human insulin and Dilp2 stimulation could selectively increase the anterograde fraction of Rab4 vesicles in *Drosophila* axons.

**Fig. 4.**
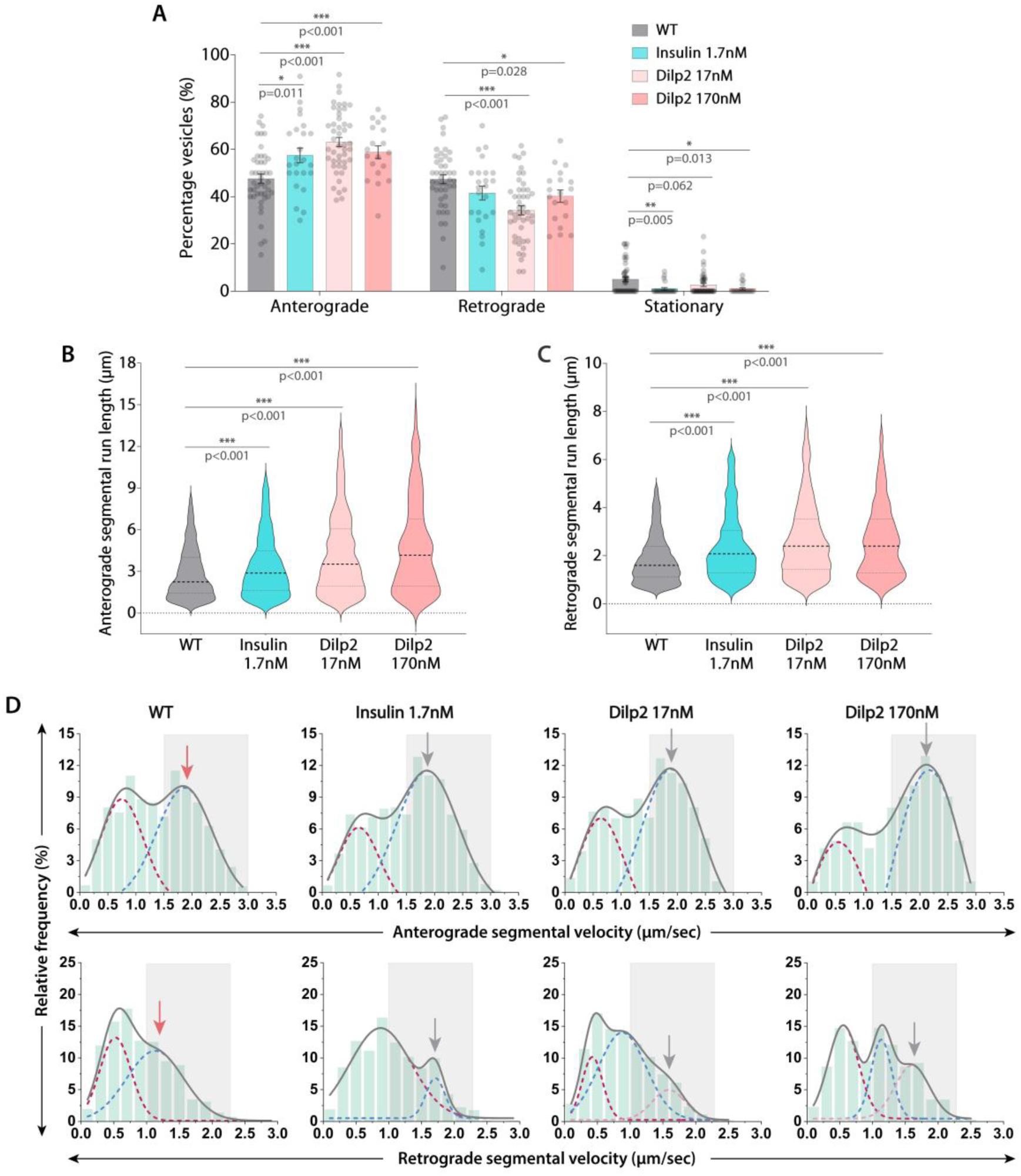
Acute stimulation with human insulin and Dilp2 increases the anterograde fraction, velocity, and run length of Rab4 vesicles in axons. **(A)** Relative distribution of the Rab4 vesicle movement (n≥19 segmental nerves, N=3-5 larvae) in the wild-type control without insulin (WT) and in the presence of different concentrations of insulin and *Drosophila* insulin-like peptide (Dilp2) at 90 hours AEL. **(B, C)** The anterograde (B) and retrograde (C) segmental run length (µm) of Rab4 vesicles in the wild-type control (WT) and different treatment backgrounds (n>300 runs) at 90 hours AEL. The pairwise significance of difference was estimated using the Mann-Whitney U-test. **(D)** Anterograde (top row) and retrograde (bottom row) segmental velocity of Rab4 vesicles in the wild-type control (WT) and different treatment backgrounds at 90 hours AEL. The cumulative distribution (grey) as a sum of multiple Gaussians to highlight slow (maroon) and fast-moving (blue and pink) populations. Grey box marks the fast-moving runs (≥1.5 µm/sec for anterograde and ≥1.0 µm/sec for retrograde).

Detailed analysis of the movements suggested that Dilp2 and human insulin treatments significantly increased the average segmental run length and frequency of fast-moving runs in the anterograde direction (Fig. 4B, D and table S1; Kolmogorov Smirnov test, p<0.001). They also increased the average segmental run length and velocity in the retrograde direction to a comparatively lesser extent (Fig. 4C-D and table S1). Though the Dilp5 treatments (both at 17nM and 170nM) did not affect the anterograde segmental run-length, the frequency of fast-moving anterograde runs was slightly reduced at higher concentrations (fig. S4B, D and table S1). Additionally, there was a moderate but significant increase in segmental run length along with a significant decrease in frequency of fast-moving runs in the retrograde direction (fig S4C-D and table S1; Kolmogorov Smirnov test, p<0.001).

Together, these observations established that acute stimulation with human insulin and Dilp2 can preferentially enhance the anterograde motility of a subset of Rab4 vesicles in a manner similar to that of constitutive InR activation in cholinergic neurons. The effects occurs at a relatively fast time scale and human insulin was more effective owing likely to its enhanced stability [29,30]. Thus, this system allowed to us further probe the fast-acting downstream pathways.

### Vps34 activity is required for stimulating the anterograde movement of Rab4 vesicles downstream of insulin signaling

Type I Phosphatidylinositol-3-Kinases (PI3Ks) was shown to regulate Rab4 activation and endosomal turnover in adipocytes and neurons [8,18,31]. Also, pan-PI3K inhibitor treatment blocked the InR^CA^-dependent enhancement of the anterograde transport of Rab4 vesicles (Fig. 3A), indicating involvement of phosphoinositide-mediated lipid signaling in the process. *Drosophila* sensory neurons express three different classes of PI3Ks [24,32], each with distinct subcellular localization pattern [33]. To identify the specific class of PI3K involved in the activation of anterograde transport of Rab4 vesicles downstream to insulin signaling, we conducted a combinatorial screen using *in vivo* live imaging of Rab4 vesicles in the presence of class-specific chemical inhibitors and tissue-specific knockdown of all three PI3Ks (movies S4-S5).

Acute inhibition (15 minutes) of all classes of PI3Ks using a pan-PI3K inhibitor (LY294002) significantly reduced the anterograde fraction (∼14%), average segmental run length, and the frequency of fast-moving anterograde runs (∼22%; Fig. 5A-C and table S1; Kolmogorov Smirnov test, p<0.001) of Rab4 vesicles. Likewise, acute inhibition of Class III PI3K (Vps34) using SAR405 led to a similar effect (Figure 5A-C and table S1), thereby indicating a possibly direct role of Vps34 in this process. In contrast, acute inhibition of the Class-I PI3K using HS173 did not affect the anterograde fraction (Fig. 5A), although it significantly altered the frequency of fast-moving runs (Fig. 5C and table S1; Kolmogorov Smirnov test, p<0.01). Consistent with the effect of insulin treatment and perturbations of insulin signaling (Fig. 4 and fig. S3), we observed relatively smaller change in the retrograde transport parameters after PI3K inhibitor treatments (fig. S5).

**Fig. 5.**
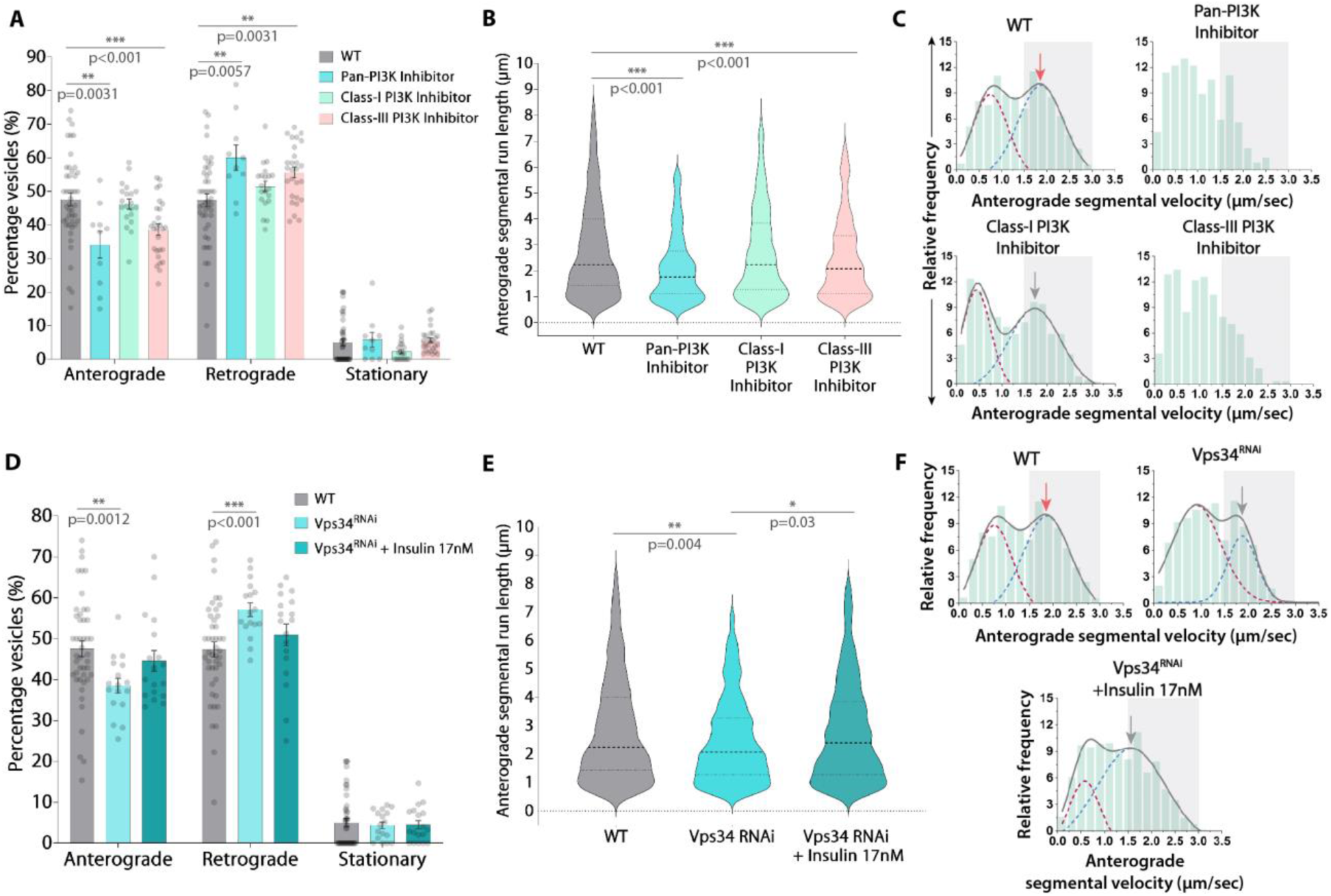
Acute inhibition and knockdown of Class-III PI3K/Vps34 reduces anterograde fraction, run length, and velocity of Rab4 vesicles in axon. (A-C) Relative distribution (A, n>10 segmental nerves, N=3-5 larvae), anterograde segmental run length (B, n>800 runs) and anterograde segmental velocity distributions (C) of Rab4 vesicles in the wild-type control (WT) and in the presence of different class-specific PI3K inhibitors at 90 hours AEL. The cumulative distribution (grey) as a sum of two Gaussians highlights slow (maroon) and fast-moving (blue) populations. Grey box marks the fast-moving (≥ 1.5 µm/sec) runs. **(D-F)** The relative distribution of the Rab4 vesicle movement (D, n≥17 segmental nerves, N=3-5 larvae), anterograde segmental run length (E, n>700 runs), anterograde segmental velocity distributions (F) in the wild-type control (WT) and Vps34^RNAi^ background in the absence and presence of insulin at 90 hours AEL. The cumulative distribution (grey) as a sum of two Gaussians highlights slow (maroon) and fast-moving (blue) populations and the grey box marks the fast-moving (≥ 1.5 µm/sec) runs. The pairwise significance of difference was estimated using the Mann-Whitney U-test.

Since Class-II PI3K inhibitor is not commercially available and the effects of Vps34 inhibitor treatment largely phenocopied the effects observed with pan-PI3K inhibitor treatment, these findings suggest that Vps34 activity could selectively regulate the anterograde fraction of Rab4 vesicles in axons.

To confirm the conjecture, we individually knocked down each one of the three PI3Ks in cholinergic neurons using established UAS-RNAi lines. This revealed that only Vps34 RNAi could significantly reduce the anterograde fraction by nearly 10% (Fig. 5D and fig. S6), as well as the run length (Fig. 5E), and the frequency of fast-moving runs (∼11%; Fig. 5F and table S1; Kolmogorov Smirnov test, p<0.001) of Rab4 vesicles at 90 hours AEL. Furthermore, acute insulin stimulation in the Vps34 RNAi background failed to increase the anterograde fraction of Rab4 vesicles to the expected level (Fig. 5D) and only marginally improved the run length (Fig. 5E) and frequency of fast-moving anterograde runs (∼4%; Fig. 5F).

On the other hand, knockdown of the PI3KC1-catalytic subunit had no significant effect on the anterograde fraction, segmental run length and the frequency of fast-moving anterograde runs (fig. S6A, B, and D). Though knockdown of the PI3KC1-regulatory subunit marginally decreased the frequency of fast-moving anterograde runs and significantly increased both the anterograde and retrograde run length (fig. S6), it had no significant impact on the overall movements (fig. S6A). Lastly, while knockdown of Class-II PI3K significantly reduced the frequency of fast-moving anterograde runs (fig. S6 and table S1; Kolmogorov Smirnov test, p<0.001), there was a significant increase on the anterograde fraction of Rab4 vesicles in the axons (fig. S6A), contrary to what was observed with Vps34 knockdown.

In summary, the results indicate that each PI3K subtype influences the that Rab4 vesicle movements in axons with Vps34 having a direct role in accelerating the anterograde movement of a subset of these vesicles downstream of insulin signaling. This finding provided a new perspective to mechanism underlying the insulin-dependent regulation of Rab4 vesicle movements in axons.

### Vps34-dependent and PI(3)P-mediated lipid signaling can regulate the velocity of Rab4 vesicles in axons

Vps34 is an early endosome-localized [34] lipid kinase that specifically produces PI(3)P from phosphatidylinositol both *in vitro* [35] and *in vivo* [36]. Further, cellular PI(3)P can be readily detected by a genetically-encoded 2x-FYVE-GFP biosensor [37]. To identify the presence and Vps34-dependent enhancement of PI(3)P levels on motile Rab4 vesicles, we used simultaneous, dual-color, time-lapse imaging of Rab4mRFP vesicle movement in cholinergic neurons expressing Rab4-mRFP and 2x-FYVE-GFP. The PI(3)P biosensor marked the motile Rab4-mRFP vesicles in the axons in both the anterograde and retrograde directions (Fig. 6A-B and movie S6). Nearly 10% of the combined pool of 2xFYVE-GFP and Rab4 vesicles (See Methods for details) were dually marked with Rab4-mRFP and 2xFYVE-GFP (Fig. 6C, D). Acute insulin stimulation increased the fraction of the colocalized vesicles by another ∼10% (Fig. 6C, D), and the Vps34-specific SAR405 inhibitor treatment abrogated the insulin-dependent increase in the percentage of colocalized vesicles (Fig. 6C, D). These findings suggest that Vps34 activation downstream of neuronal insulin signaling could induce the PI(3)P formation on Rab4 vesicles. The formation of PI(3)P on the endosome surface activates PI(3)P-mediated lipid signaling [38]. In conjunction with the previous findings regarding the influence of InR signaling through Vps34 on Rab4 vesicle movements (table S1), the data further suggest that increased PI(3)P on the Rab4 vesicles could enhance their anterograde velocity and run-lengths.

**Fig. 6.**
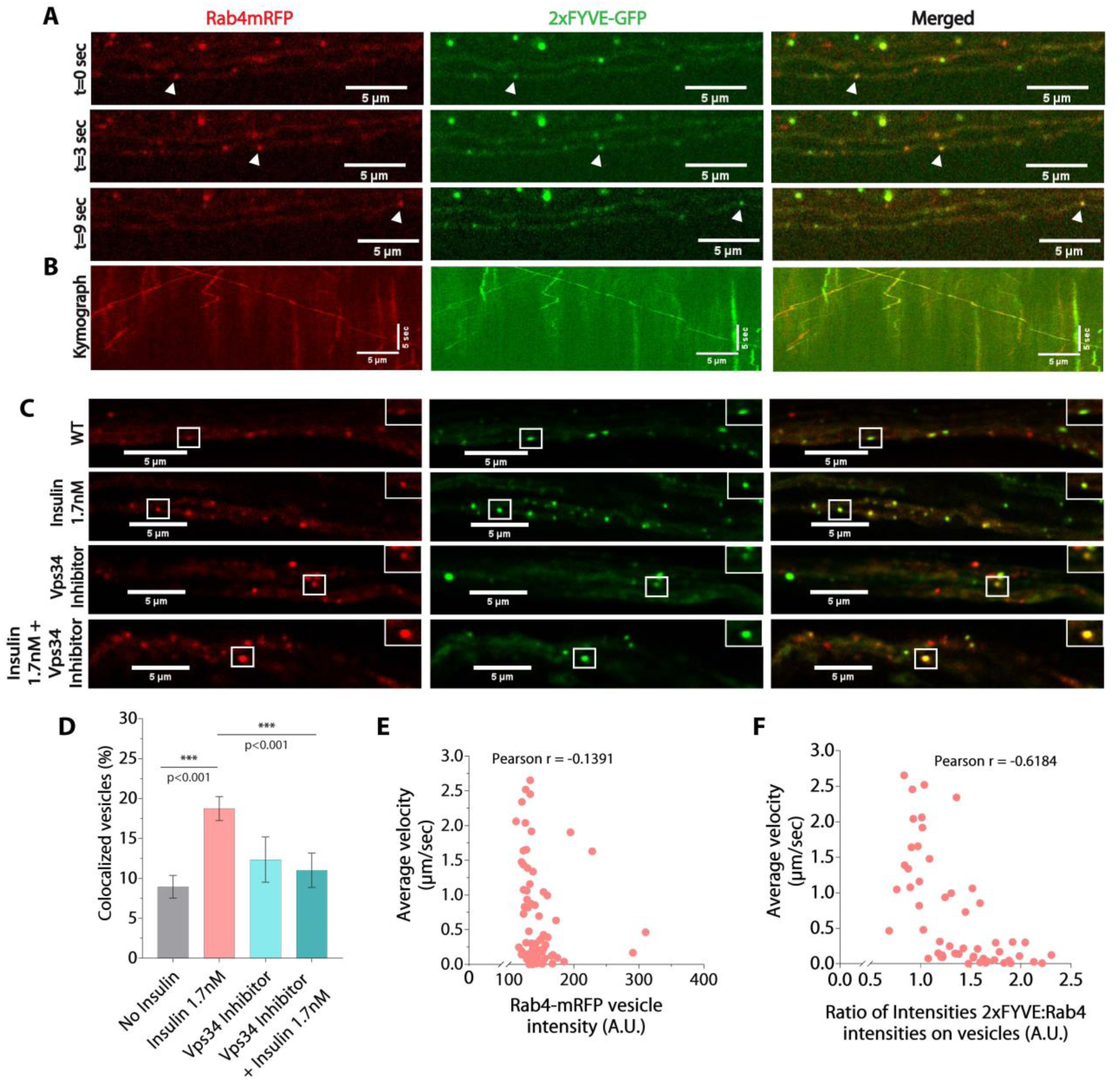
Vps34-mediated endosomal PI(3)P-signaling regulates axonal transport of Rab4 vesicles downstream of insulin signaling. (A, **B)** Simultaneously acquired dual-channel time-lapse images of segmental nerves (A) and kymographs (B) depict comigration of genetically encoded Rab4mRFP (red) and 2xFYVE-GFP (green, PI(3)P biosensor) in cholinergic axons. **(C)** Optical slices of segmental nerves of wild-type control with no insulin (WT) and after treatments with insulin and class-III PI3K inhibitor (SAR405) depict relative Rab4mRFP (red) and PI(3)P biosensor 2xFYVE-GFP (green) levels on endosomal vesicles in axons. **(D)** Percentage of colocalized vesicles (Mean ± SEM) of the total (Rab4mRFP and 2xFYVEGFP) in wild-type control and different treatment backgrounds (n≥11 ROIs, N=3 larvae, >1000 vesicles each). The pairwise significance of difference was estimated using the Mann-Whitney U-test. **(E, F)** Correlations between Rab4-mRFP intensity and corresponding average velocities obtained using single-particle tracking (E), and between the 2xFYVE-GFP:Rab4-mRFP intensity ratios and corresponding average velocities (F, n=74 vesicles, N=6 larvae) of Rab4 vesicles.

To test this hypothesis, we harnessed the previously documented masking ability of 2xFYVE-GFP [39]. A simultaneous estimation 2x-FYVE-GFP and Rab4-mRFP intensities on individual vesicles in the axons, coupled with the single-particle tracking of the same vesicles, indicated no correlation between the Rab4mRFP levels on vesicles (a proxy for Rab4 activation) and their average velocity (Fig. 6E). In contrast, increasing levels of the PI(3)P biosensor were inversely correlated with the average velocity of these vesicles (Fig. 6F), *i.e.*, the higher the intensity of the biosensor, the lower the velocity of the vesicle and *vice versa* (movie S7). Thus, abrogation of PI(3)P-mediated lipid signaling on Rab4 vesicles proportionately reduces their overall motility in the axons.

Together with the previous results, this observation further establishes that local PI(3)P-signaling on the surface of Rab4 vesicles, downstream of insulin receptor signaling, is a critical determinant of their anterograde transport properties. This effect may manifest by regulating the recruitment of the plus-end-directed kinesin motors on these vesicles. These results also suggested that PI(3)P levels on Rab4 vesicles could indicate the activation of InR signaling in axons.

### Loss of Kinesin-2 function failed to block the insulin-stimulated increase in the anterograde transport of Rab4 vesicles

Next, we aimed to identify the kinesin motor that could likely play a role in regulating axonal transport of Rab4 vesicles downstream of insulin signaling in the axons. The heterotrimeric kinesin-2 was the first candidate, as it had been established as a key regulator of Rab4 vesicle transport in cholinergic neurons [8] and that of the GLUT4-vesicles in adipocytes [18]. Using dual-channel live imaging of cholinergic neurons expressing Rab4mRFP and Klp64D-GFP (KIF3A ortholog in *Drosophila* and an essential subunit of kinesin-2), we demonstrate that kinesin-2 is likely to comigrate with Rab4 vesicles in the axons (fig. S7A-B). However, these events were infrequently detected due to very high cytoplasmic backgrounds of Klp64D-GFP (fig. S7A). We then used a previously characterized *Klp64D* mutant (*Klp64D^K5^*), known to affect kinesin-2 function *in vivo* [40], to perturb the transport of Rab4 vesicles in the axons. Consistent with the previous report in lch5 neurons of *Drosophila,* which showed a significant decrease in the anterograde fraction and segmental velocity of Rab4 vesicles in the *Klp64D^K5^* homozygous background [8], the anterograde movement of Rab4 vesicles was moderately affected in the cholinergic neurons in the heterozygous (*Klp64D^K5^/+*) background (fig. S7C-E). Additionally, we noted a significant increase in the pool of stationary vesicles (fig. S7C), suggesting a likely role of kinesin-2 in the initiation of Rab4 vesicle transport. As expected, it significantly reduced the anterograde and retrograde run lengths of Rab4 vesicles (fig. S7D-E and table S1) with a significant reduction in the frequency of fast-moving anterograde and retrograde runs (fig. S7F-G; Kolmogorov Smirnov test, p<0.001). These observations suggested that *Klp64D^K5^* mutation could act as a dominant negative allele and suppress the kinesin-2 function in transport of Rab4 vesicles. However, contrary to the expectations, acute insulin stimulation in the *Klp64D^K5^/+* background rescued the motility of the stationary pool and elevated the anterograde fraction of Rab4 vesicles to the expected level (fig. S7C). Such a rescue would ideally not be possible if kinesin-2 was the motor responsible for insulin-mediated increase in anterograde fraction of Rab4 vesicles. Hence, these results suggest that while kinesin-2 could play a critical role in promoting the Rab4 vesicle transport, a different anterograde motor is likely to be involved in insulin-stimulated acceleration of the subset of Rab4 vesicles in the axons.

### Klp98A recruitment via PI(3)P-signaling activates anterograde transport of Rab4 vesicles downstream of insulin stimulation

To identify this other kinesin, we focused on KIF16B, a kinesin-3 family motor that contains a PI(3)P-binding PX-domain in the C-terminal tail region, which was shown to regulate the anterograde movement of Rab5 vesicles during early endosome recycling in HeLa cells [41]. Though KIF16B was suggested to be absent from axons by a previous study [42], we tested whether Klp98A – the *Drosophila* ortholog of KIF16B – could accelerate the movement of Rab4 vesicles downstream to insulin signaling in the axons. First, we demonstrated that ectopically expressed Klp98A-GFP comigrated with Rab4-mRFP vesicles in the axons (Fig. 7A and movie S8). Next, cell-specific Klp98A knockdown significantly reduced the anterograde fraction, run length, and segmental velocity of Rab4 vesicles (Fig. 7C-E, table S1). As observed previously, it also led to a significant reduction of retrograde segmental run length and a moderate reduction in retrograde velocity (fig. S8A-B), which was much less severe than the loss of anterograde transport parameters. Finally, acute insulin stimulation did not increase the anterograde fraction, run length, and velocity of Rab4 vesicles in the Klp98A^RNAi^ background (Fig. 7C-E). Altogether, these observations suggested that insulin stimulation could help recruit Klp98A onto the Rab4 vesicles through Vps34/PI(3)P signaling.

**Fig. 7.**
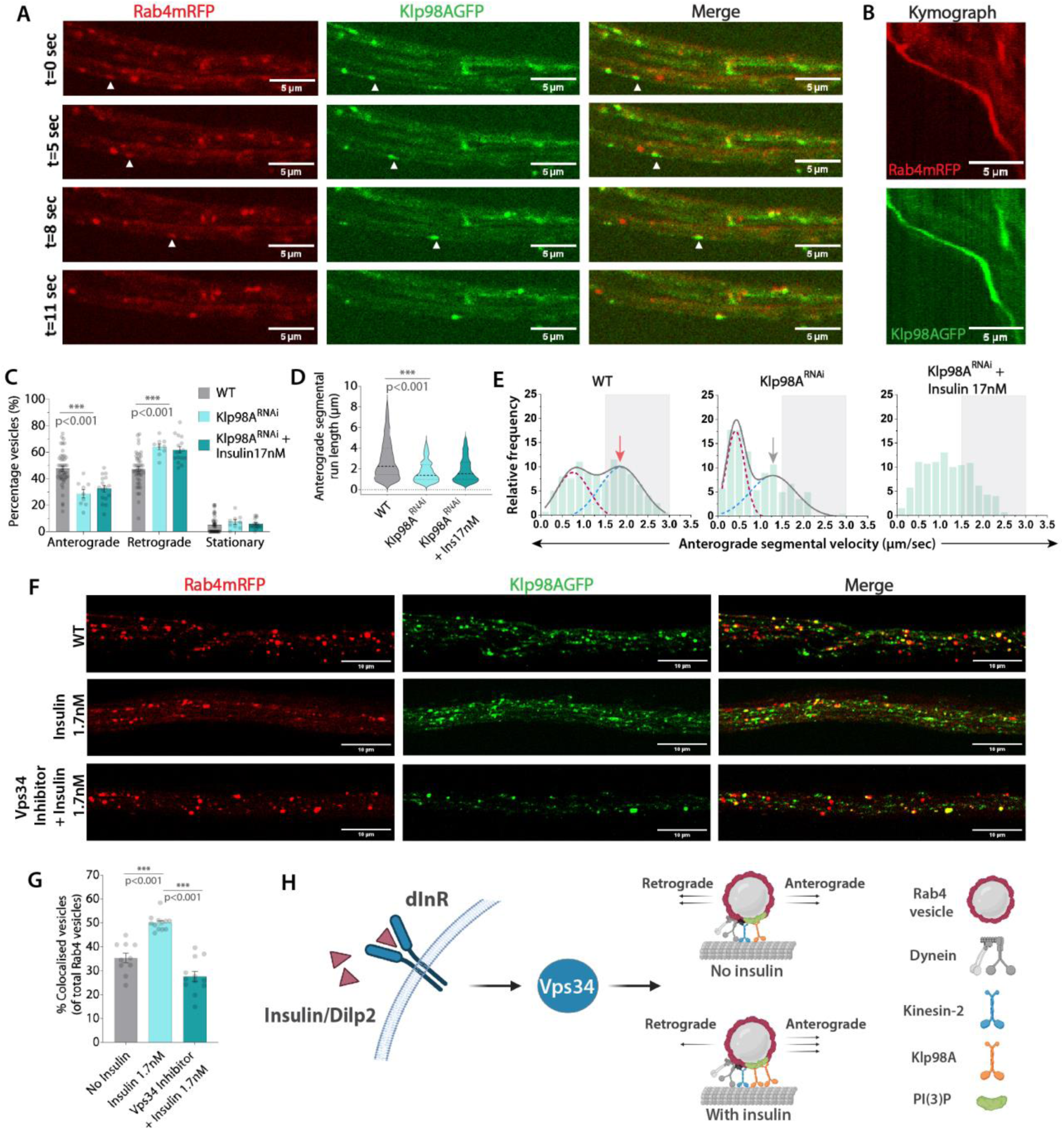
Klp98A (KIF16B ortholog) recruitment through a Vps34/PI(3)P-dependent pathway downstream of insulin signaling increases the anterograde fraction of Rab4 vesicles. (A, **B)** Dual-channel time-lapse images of segmental nerve (A) and kymographs (B) depict comigration of genetically encoded Rab4mRFP and Klp98AGFP in cholinergic axons. **(C-E)** Relative distributions of the Rab4 vesicle movement (C, n≥9 segmental nerves, N=4-5 larvae), anterograde segmental run lengths (D) and anterograde segmental velocity distributions (E, n>200 runs) in the wild-type control (WT) and Klp98A^RNAi^ backgrounds in the absence and presence of insulin at 90 hours AEL. The cumulative velocity distribution (grey) as a sum of two Gaussians highlights slow (maroon) and fast-moving (blue) populations. **(F)** Optical slices of segmental nerve axons in control (WT), and after acute treatment with insulin, and insulin+Vps34 inhibitor depict the colocalization of Rab4mRFP and Klp98AGFP. **(G)** Fraction (Mean+S.E.M) of Rab4mRFP and Klp98AGFP colocalized vesicles in different treatment backgrounds (n≥10 segmental nerves, N≥3 larvae, ≥1800 vesicles). The pairwise significance of difference was estimated using the two-tailed t-test. **(H)** Schematic depicting a summary of signaling cascade where activation of insulin signaling via Dilp2-InR-Vps34 axis recruits the PI(3)P-binding kinesin-3 motor Klp98A on Rab4 vesicles and increases their anterograde fraction in the axons.

To test this hypothesis, we estimated the number of Rab4-mRFP vesicles labelled with the Klp98A-GFP in segmental nerve axons in the presence and absence of insulin. It revealed that acute insulin stimulation could significantly increase the frequency of Klp98A-GFP localization on Rab4 vesicles and this increase was completely abrogated in the presence of Vps34-specific inhibitor (Fig. 7F-G; n≥10 segmental nerves, N≥3 larvae, >1800 total vesicles). Molecularly, our data demonstrated that Klp98A recruitment on Rab4 vesicles could be regulated by the insulin-dependent activation of Vps34 (Fig. 7H), which would be a critical step in increasing the anterograde velocity, processivity and fraction of Rab4 vesicles in the axons. These results implicated Klp98A/KIF16B in accelerating the Rab4 vesicles towards synapse upon insulin stimulation. Additionally, our results also suggested that kinesin-2 could play an important role in initiating anterograde movement of Rab4 vesicles (fig. S7C). In effect, a concerted action of these two kinesin motors in the axon could propel the Rab4 vesicles towards the synapse. Lastly, given the previous reports [8,18], our data also implicated a crucial role of Rab4 activity in recruiting both these kinesin motors on endosomal vesicles.

### PI(3)P-signaling and Klp98A recruitment on Rab4 vesicles is developmentally regulated

The results described so far clearly establish that the InR-Vps34-Klp98A axis could accelerate the synapse-bound movements of a subset of Rab4 vesicles marked with

PI(3)P. Also, the presence of PI(3)P or Klp98A-GFP on the Rab4 vesicles effectively reported the InR activation in axons. Hence, to understand whether this pathway could indeed account for the programmed changes in the Rab4 vesicle movement from 72-90 hours AEL, we studied the developmental changes in the 2xFYVE-GFP and Klp98A-GFP levels on Rab4 vesicles in axons using fixed tissue preparations of transgenic larvae at 72, 80, and 90 hours AEL.

First, we observed a significant increase in the colocalization of Rab4-mRFP and 2xFYVE-GFP during 72-80 hours AEL, which subsided at 90 hours AEL (Fig. 8C-D), indicating a developmentally regulated increase of the InR signaling in cholinergic neurons at 80 hours AEL. A similar episodic increase in the fraction of vesicles marked with both Rab4-mRFP and Klp98A-GFP occurred at 80 hours AEL and subsided at 90 hours AEL (Fig. 8A-B). The pattern of Klp98A-GFP recruitment is consistent with the trend observed for the anterograde fraction of Rab4 vesicles during these periods (Fig. 2D), indicating that Klp98A-GFP levels on Rab4-mRFP vesicles could serve as a determinant for regulating anterograde transport of Rab4 vesicles.

**Fig. 8.**
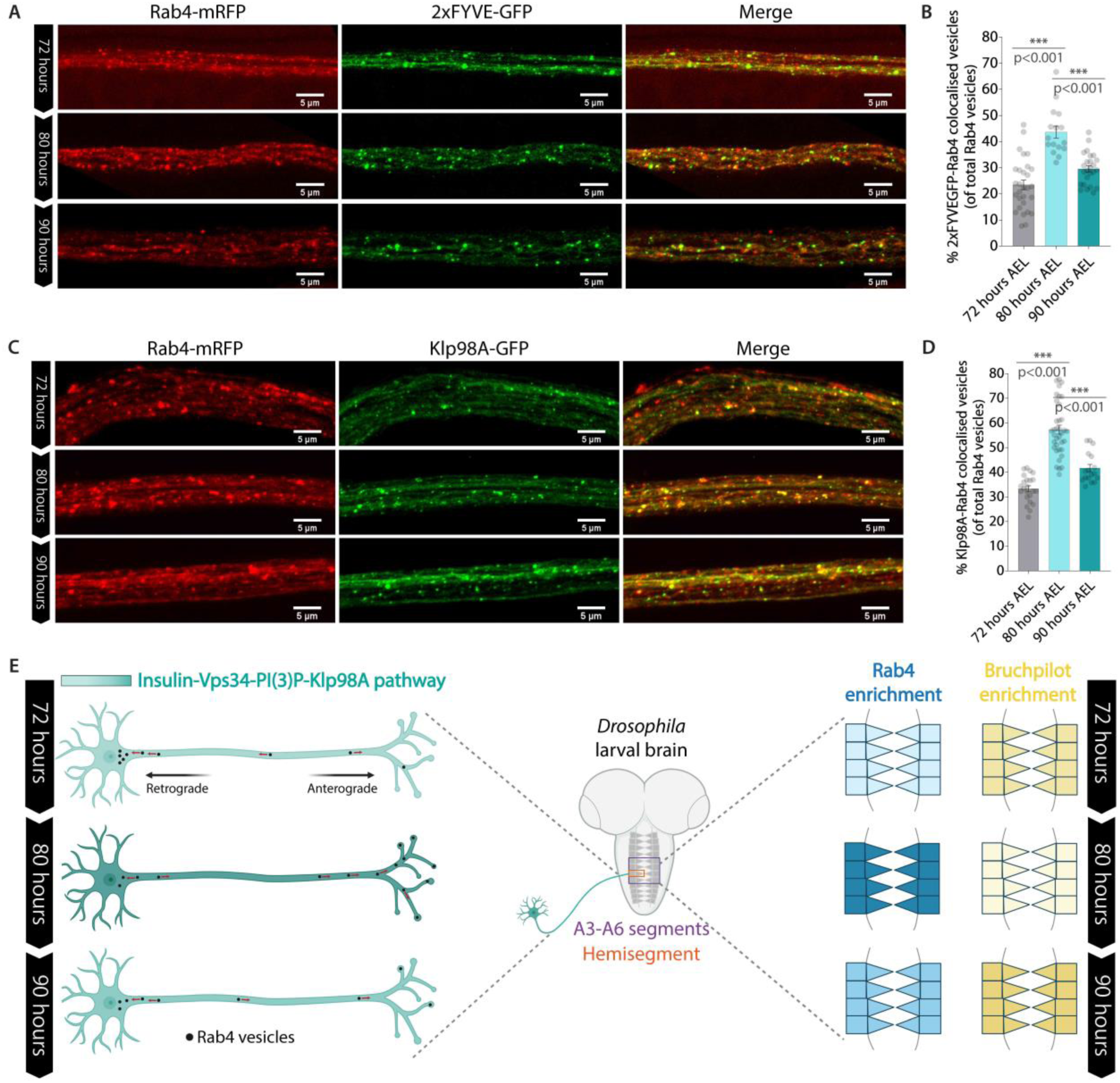
Klp98A and PI(3)P levels on Rab4 vesicles are developmentally regulated in sync with each other. **(A)** Optical slices of segmental nerve axons depicting colocalization between Rab4 vesicles (red) and 2xFYVE-GFP, PI(3)P biosensor (green) at 72-90 hours AEL. **(B)** Percentage (Mean ± SEM) of Rab4 vesicles colocalized with PI(3)P biosensor at 72-90 hours AEL (n>16 segmental nerves, N=3-6 larvae). The pairwise significance of the difference is estimated using the two-tailed t-test. **(C)** Optical slices of segmental nerve axons depicting colocalization between Rab4 vesicles (red) and Klp98A motor (green) at 72-90 hours AEL. **(D)** Percentage (Mean ± SEM) of Rab4 vesicles colocalized with Klp98A motor at 72-90 hours AEL (n>17 segmental nerves, N=3-6 larvae). The pairwise significance of the difference is estimated using the two-tailed t-test. **(E)** Working model shows that developmentally programmed insulin signaling in the neurons activates Vps34, a lipid kinase that produces PI(3)P on Rab4 vesicles. Increased levels of PI(3)P on Rab4 vesicles recruit the fast-moving anterograde motor Klp98A that elevates their anterograde fraction in the axons, increase its presynaptic enrichment, and promotes synaptic remodeling in the CNS.

Together, these two findings suggest that that developmentally programmed insulin signaling would operate through Vps34 activation and regulate the recruitment of Klp98A on Rab4 vesicles via PI(3)P. The recruitment of a fast kinesin motor would stimulate the anterograde movement of these vesicles resulting in the presynaptic enrichment of Rab4, which is likely to induce neurite growth and synaptic remodeling in *Drosophila* larval CNS.

## Discussion

### Effects of neuronal insulin signaling on Rab4-associated recycling endosomes in axons

Endosomal trafficking and recycling in neurons are essential for neuronal development and survival [43,44]. The directed movement of endosomal traffic is a critical regulator of several neuronal processes like signaling, autophagy, synaptic vesicle recycling, and neurotransmission [45–47]. Besides, perturbations in endosomal functions are one of the earliest pathologies in neurodegenerative disorders like Alzheimer’s disease [48]. However, signaling cues regulating directed endosomal movement in the axons remain relatively understudied. In this context, we show that developmental regulation of neuronal insulin signaling could regulate the directionality and overall flow of a subset of endosomal vesicles in axons.

Insulin signaling in *Drosophila* brain has been implicated in age-related cognitive decline [14]. Besides, insulin signaling in neurons has been shown to regulate synaptic density [49], neurodegeneration [50], aging [51], and behavior [52] in various other model systems. However, its effect on long-range axonal transport, which is essential for maintenance of synaptic homeostasis, remained unknown. Using ectopic stimulation and tissue-specific genetic perturbations, we show that signaling via insulin/Dilp2 and dInR could accelerate the anterograde movement of a subset of Rab4 vesicles in the cholinergic neurons of *Drosophila*. This observation could have substantial physiological bearings as increased accumulation of activated Rab4 at the presynaptic terminals is known to adversely affect synaptic density in *Drosophila* CNS [8], and elevated Rab4 levels in both CBF and hippocampal CA1 neurons are positively correlated with neurodegeneration and cognitive decline in Alzheimer’s disease [9,10]. Our results, together with other findings suggest that insulin signaling could manifest its effect on synapse organization, neurodegeneration, and aging, in part via altered trafficking of Rab4 endosomes.

The experimental data also suggest that insulin-dependent redirection of Rab4 may regulate neuropil growth and synaptic remodeling in the larval VNC. Rab4 plays an integral role in neurite outgrowth, recycling of surface receptors, and regulating endosomal traffic inside a cell [1–3,53]. Thus, increased enrichment of Rab4 at the presynaptic terminals could alter recycling of synaptic vesicles and promote neurite outgrowth, consequently regulating synaptic strength and stability. Such a process may also modify the composition of presynaptic membrane and alter neurotransmission.

### Insulin signaling preferentially activates Vps34, increasing PI(3)P on Rab4 vesicles in axons

This study also identified a selective activation of Vps34 downstream of insulin signaling in the axons. Insulin signaling is known to activate PI3Kinases, particularly the Class I PI3K in peripheral tissues [54], which activates Rab4 and early endosomal recycling [31]. Here, we show that insulin stimulation preferentially activates Class-III PI3K, also known as Vps34, which is involved in accelerating the speed of Rab4 vesicles. Thus, the mechanism activating the Rab4 vesicle trafficking downstream of the insulin receptor appears to be quite distinct in the axons as compared to peripheral tissues. Vps34 – the lipid kinase that exclusively produces PI(3)P inside the cell – plays a key role in regulating neuronal processes like autophagy [55], synaptic vesicle recycling [56], neurotransmission [57], and neuronal survival [58]. Recent studies have also shown that PI(3)P-positive endosomes have a critical role in regulating synaptic vesicle recycling and neurotransmission [45]. Further, a nutrient-sensitive, PI(3)P-mediated lipid signaling on endosomes was suggested to regulate ER shape and mitochondrial function [59]. However, the source, identity, and mechanism(s) regulating PI(3)P formation on endosomes in these studies was unclear. Our study shows that neuronal insulin signaling promotes Vps34-dependent activation of anterograde transport of Rab4-PI(3)P vesicles, which could also act as an active source of replenishing the pool of PI(3)P-Rab4 dual-positive vesicles enriched at the synapses.

### Klp98A recruitment and selective augmentation of synapse-directed Rab4 vesicle movement in axon

Finally, the data suggest that the Rab4-PI(3)P vesicles are accelerated by recruiting a specific type of kinesin-3 motor, Klp98A, which is the *Drosophila* ortholog of mammalian KIF16B. These motors contains a PI(3)P-binding PX-domain in the C-terminal tail [60]. Klp98A recruitment onto Rab4 vesicles in the axons due to Vps34 activation downstream of acute insulin stimulation increases their speed by nearly 2-folds. An unbiased proteomic screen to identify all Rab4 effectors identified Vps34 as one of the Rab4 effectors [61], suggesting a potential recruitment mechanism for Vps34 on Rab4 vesicles for the local production of PI(3)P. The results described here linked Vps34 activity to Klp98A/KIF16B recruitment on Rab4 vesicles and suggested that the Vps34 activation downstream of insulin signaling in the axon could critically determine the quantity of Rab4 vesicle movement towards the synapse.

A previous study in mouse hippocampal neurons showed that KIF16B is required for somatodendritic localization of early endosomes which helps in the trafficking of AMPA and NGF receptors [42]. However, whether there is any role of KIF16B in regulating long-range axonal transport *in vivo* was unclear. We also noted that not all motile Klp98A-positive vesicles in axons have Rab4, indicating that Klp98A transports other cargos apart from Rab4 vesicles in axons. Interestingly, mutations in the cargo-binding PX domain of KIF16B have recently been reported in patients with intellectual disability syndrome [62], although the underlying cause remains to be investigated. Thus, our results could provide a critical mechanistic input because perturbations in Rab4 levels and functionality also affect synaptic homeostasis, which happens to be one of the hallmarks of intellectual disabilities [63]. Therefore, it will also be worth investigating in the future if there is a synergistic interaction between Rab4 and Klp98A keeping in mind their overlapping functions and phenotypes in the CNS.

Finally, cell biological studies suggested that KIF16B drives the fission of early endosomes to form tubules [64], receptor recycling and degradation [41], transcytosis [65], and polarized transport of growth factor receptors [60]. In addition, the Klp98A motor is implicated in the maturation of autophagic vesicles by promoting fusion through a motor-independent function [66]. The formation of PI(3)P on Rab11 vesicles upon starvation has been shown to serve as a platform for autophagosome formation in HeLa cells [67]. However, such platforms and/or the identity of PI(3)P membranes required for autophagosome formation in neurons were not identified. In this context, it may be conjectured that the activation of Vps34-PI3P-Klp98A-dependent movement of Rab4 vesicles towards the synapse in *Drosophila*, as revealed by the above data, could serve as a platform for regulating autophagosome formation at the synapse and reduce the overall synaptic volume. This supposition is also consistent with the results indicating an inverse correlation between Rab4 enrichment at the presynaptic terminals and synaptic density in a developing *Drosophila* larva.

## Material and Methods

### Drosophila stocks and rearing

All fly stocks were obtained from Bloomington Drosophila stock center unless mentioned otherwise and were reared at 25°C on standard corn agar meal with a 12-hour light-dark cycle. Eggs were collected for 1 hour and kept at 25°C for 72, 80, or 90 hours. Fly stocks used: Brp-GFP (BL59292), *UAS Rab4mRFP* (BL8505), *UAS dsRNA InR* (BL31594, Valium1), *UAS dsRNA InR* (BL51518, Valium20), *UAS InR^CA^* (BL8250), *UAS dsRNA PI3K92E* (BL27690), *UAS dsRNA PI3K21B* (BL38991), *UAS dsRNA PI3K68D* (BL35265), *UAS dsRNA PI3K59F* (BL33384), *UAS GFP-myc-2xFYVE* (BL42712), *Klp64D^K5^* [40] and *UAS Klp64D-GFP* [68], *UAS dsRNA Klp98A* (BL 50542), and *UAS Klp98A-GFP* [69].

### Larval fillet preparation: Time-lapse imaging and fixed tissue preparations

Larvae were dissected in 1x HL3.1 buffer [70] (containing 70 mM NaCl, 5 mM KCl, 1.5 mM CaCl2, 4 mM MgCl2, 10 mM NaHCO3, 5 mM Trehalose, 115 mM Sucrose, and 5 mM HEPES) and insulin (Sigma-Aldrich), Dilps (Pheonix Pharmaceuticals Inc.), PI3K inhibitors (Sigma-Aldrich) were finally reconstituted in HL3.1 buffer for treatment followed by mounting on the coverslip cavity chamber. A small piece of tissue drenched in water was placed on the side of the petri-dish to maintain humidity. The dish is then covered and sealed with Parafilm and immediately taken for time-lapse imaging. Epifluorescence and spinning disc confocal (SDC) time-lapse imaging was performed using Nikon Ti Eclipse TIRF Microscope (installed with a Yokogawa CSU-W1 disk with pinhole size 50 µm for SDC microscopy) operated with running Nikon Elements software using a 100x (1.4 NA) oil-immersion objective (binning=1) with Andor iXon EMCCD camera and sCMOS camera (Zyla 4.2 plus sCMOS) respectively.

Live movies were recorded for all conditions at a frame rate of 8-10 frames per second (fps). For fix tissue analysis, fillet preparations were immediately fixed in 4% paraformaldehyde in 1X PBS for 15-20 minutes at room temperature (RT). Following fixation, they were rinsed thrice with 1X PBS and mounted on a coverslip with a drop of VECTASHIELD® antifade mounting medium (Vector laboratories).

### Insulin/Dilp and drug treatments

Stock solutions of Insulin (Sigma, I0516) and Dilp2/5 (Pheonix Pharmaceuticals Inc., 036-17 & 035-96) were diluted and prepared in HL3.1 buffer, respectively. Likewise, stock solutions of non-specific PI3K inhibitor (5mM) LY294002 (Abcam, #ab120243), HS173 (50µM) (Sigma Aldrich, 5.32384), SAR405 (5mM) (Sigma-Aldrich, 5.33063) were prepared in DMSO (Sigma) and used at effective concentrations of 50 µM, 50 nM, and 25 µM reconstituted in HL3.1 buffer respectively.

### Quantification of axonal transport and statistical analyses

All live-imaging movies were analysed on ImageJ/Fiji® (https://imagej.net/Fiji). KymoAnalyzer plugin [71] was used for estimation of axonal transport parameters like velocity, run length, and cargo population. Input pixel size and frame rate were used as per the movies recorded.

### Immunostaining

Dissected VNC of larvae were immediately fixed in 4% paraformaldehyde (PFA) in 1x PBS for 20 minutes at room temperature followed by 3 washes with 1x PBS, 10 min each. These samples were then permeabilized in 1x PBS containing 0.3% Triton-X-100 (PTX) for 20 minutes followed by blocking for an hour with 1 mg/ml Bovine Serum Albumin (BSA) in 0.3% PTX (PBTX) at room temperature to block the non-specific reactive binding sites for antibody. Samples were then incubated with the primary antibodies diluted in PBTX for 2 hours at room temperature followed by three 10-minute washes in PBTX. Incubation with secondary antibodies diluted 1:400 in PBTX was done for an hour followed by three 10-minute washes in PBTX. Samples were finally mounted on a glass slide in a drop of Vectashield® (Vector Laboratories Inc., USA) under an 18 mm X 18 mm coverslip of 0.17 mm thickness. Same procedure was followed for immunostaining of fillet preparations. Antibodies, stains, and dyes used: anti:Brp (nc82-DSHB, 1:200), anti-GFP (ab290-Abcam, 1:1000), anti-Rab4 (ab78790-Abcam, 1:400), Alexa Fluor 488 (A11029, A11008 – Invitrogen, 1:400), and Hoechst 33342 (Sigma Aldrich, 1:100).

**3D rendering: volume and intensity estimation using Imaris® software** All fluorescence images of Bruchpilot immunostained larval VNC were collected under constant acquisition conditions using Zeiss LSM 510 Meta laser scanning confocal microscope, using a 40x /1.3 NA objective at a pixel resolution of 0.62 X 0.62 μm^2^. Same secondary antibody was used across various time points and genotypes. The acquisition parameters viz., laser power, PMT gain, scan speed, optical zoom, offset, step size, pinhole diameter was kept constant for each experimental data set and samples were processed in a single batch. Volume and total intensity measurements on 3D volume rendered image stacks were done using ImageJ® (https://imagej.nih.gov/ij/) and Imaris® software. A surface was reconstructed on the 3D volume rendered image using the surface module of the object menu of Imaris software for each neuromere hemisegment by manually selecting the regions. Intensity and volume measurements were taken from these surfaces.

### Electron microscopy

VNCs from larvae at 72, 80, and 90 hours AEL were dissected in HL3.1 buffer and fixed overnight in 2.5% glutaraldehyde (EM Sciences), 4% paraformaldehyde, and 0.04% CaCl2 in 0.1 M phosphate buffer at 4°C. The tissues were washed in 0.1 M phosphate buffer and post-fixed in OsO4 for 4 hour at 4°C, followed by washes in 0.1 M phosphate buffer (pH 7.4), dehydration in graded series of ethanol, and embedding in Araldite (Merck). Ultrathin sections were obtained in Leica EM UC6, stained with aqueous uranyl acetate and lead citrate, and imaged using a Zeiss Libra 120 EFTEM.

### Colocalization estimation

Percentage of colocalized vesicles (Fig 6-8) were estimated using a standard automated ImageJ plugin ComDet v0.5.5 developed by Eugene Katrukha from Utrecht University https://github.com/UU-cellbiology/ComDet). Imaging parameters for ComDet were chosen such that we could detect >95% vesicles in each image (adjusting pixel size, intensity threshold, and distance between colocalised spots) and automated analysis was validated using manual counting for the control dataset.

### Statistical information

Origin™ 2020 was used for plotting the frequency histograms (bin size 0.2 µm/sec) and curve fitting was done using Multiple Peak Fit with Gaussian Peak Function. GraphPad Prism 9.5.1 was used for plotting scatter plots and calculation of the Pearson coefficient (r). Nonparametric Kolmogorov-Smirnov test for comparing all the frequency histograms for segmental velocities and was performed using Origin™ 2020. Mann-Whitney U test and Student’s t-test were performed using GraphPad Prism 9.5.1. The statistical tests used, ‘N’, and ‘n’ values are specified in the main text/figure legends for all the figures.

## Data availability statement

All data are available in the main text or the supplementary materials.

## Acknowledgements

We thank all K.R. and Howard lab members for all the help and technical support. We also thank Sundar Ram Naganathan for comments on an earlier version of the manuscript. We acknowledge Hugo Stocker for sharing InR-CFP and Emmanuel Derivery for sharing Klp98A fly stocks; Swagata Dey and Komal Raina for discussions. We acknowledge the use of Biorender for making schematics and Adobe Illustrator for making the figures.

## Author Contributions

Conceptualization: KS, SD, DR, KR; Investigation: KS, SD, DR, AS; Methodology: KS, DR, KR; Visualization: KS, SD, DR, KR; Formal analysis and Validation: KS, SD, DR, AS, SS, JH, KR; Data Curation: KS, SD, DR; Writing – Original Draft preparation: KS, KR; Writing – Review and Editing: KS, SD, DR, AS, SS, JH, KR; Funding acquisition: KS, JH, KR; Supervision: KR, JH (at Yale).

