## Supplemental Figures for "PI(3)P signaling regulates endosomal flux underlying developmental synaptic remodeling via Rab4"

**The PDF file includes:**

Materials and Methods
Supplementary Text
Figs. S1 to S8
Table S1
References

**Other Supplementary Materials for this manuscript include the following:**

Video S1 to S8
Data S1 to S10

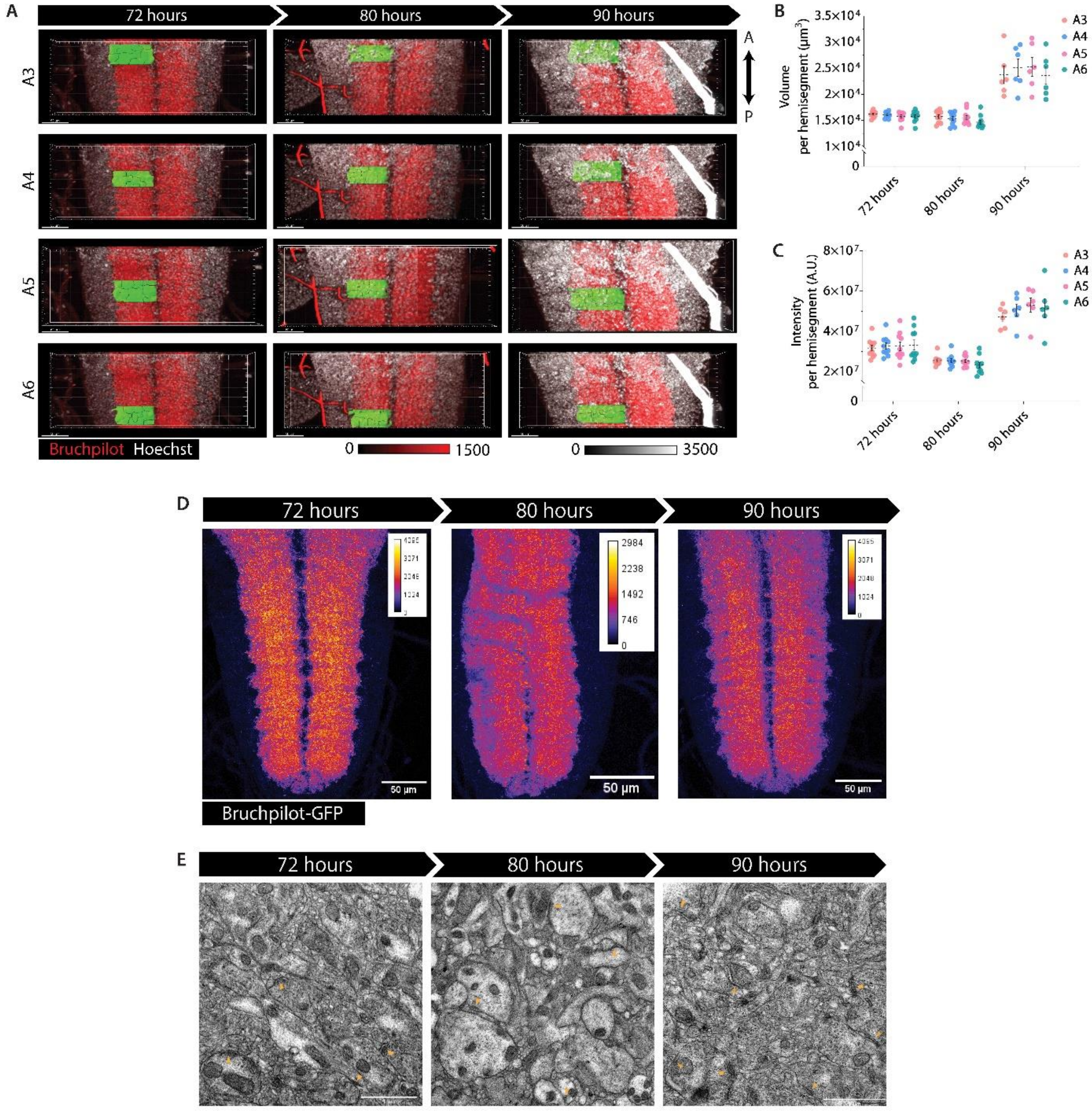


**Supplementary Figure 1: The Drosophila central nervous system (CNS) undergoes extensive synaptic remodelling during the 3rd Instar larva stage.** (A) Maximum intensity projections of a larval VNC marked with endogenous Bruchpilot-GFP (MiMIC fly line #BL59292) from 72-90 hours AEL. (B) A3-A6 segments of larval VNC stained with anti-Bruchpilot (red) and Hoechst (white) at 72-90 hours AEL. (C, D) Mean ± SEM of synaptic volumes (C) and Bruchpilot enrichment (D) in A3-A6 hemisegments (N=3-5 larvae). (E) Electron micrographs of larval VNC at 72 (left), 80 (middle), and 90 hours (right) AEL where yellow arrowheads highlight active zones in synaptic boutons. Note the apparent increase in their number and electron density of active zones at 90 hours AEL (right), in agreement with Bruchpilot intensity and volume measurements.

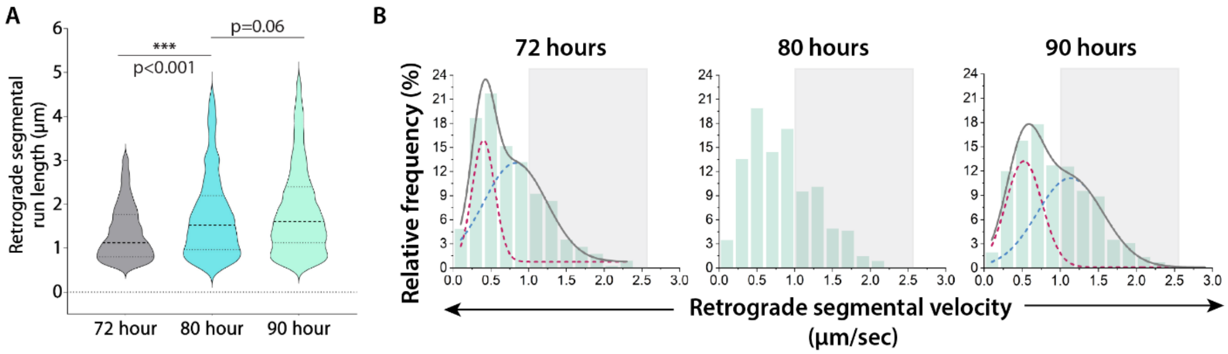

**Supplementary Figure 2: Retrograde axonal transport of Rab4 vesicles in a developing larva from 72-90 hours AEL.** (A) Retrograde segmental run length of Rab4 vesicles at 72-90 hours AEL (n>300 runs from N=3-5 larvae). The pairwise significance of difference was estimated using the Mann-Whitney U-test. (B) Retrograde segmental velocity distributions of Rab4 vesicles at 72-90 hours AEL (n>300 runs, N=3-5 larvae). The cumulative distribution (grey) as a sum of two Gaussians (maroon and blue dotted lines) highlights slow (maroon) and fast-moving (blue) populations (See methods for details). Grey box marks the fast-moving (≥ 1.5 µm/sec) runs.

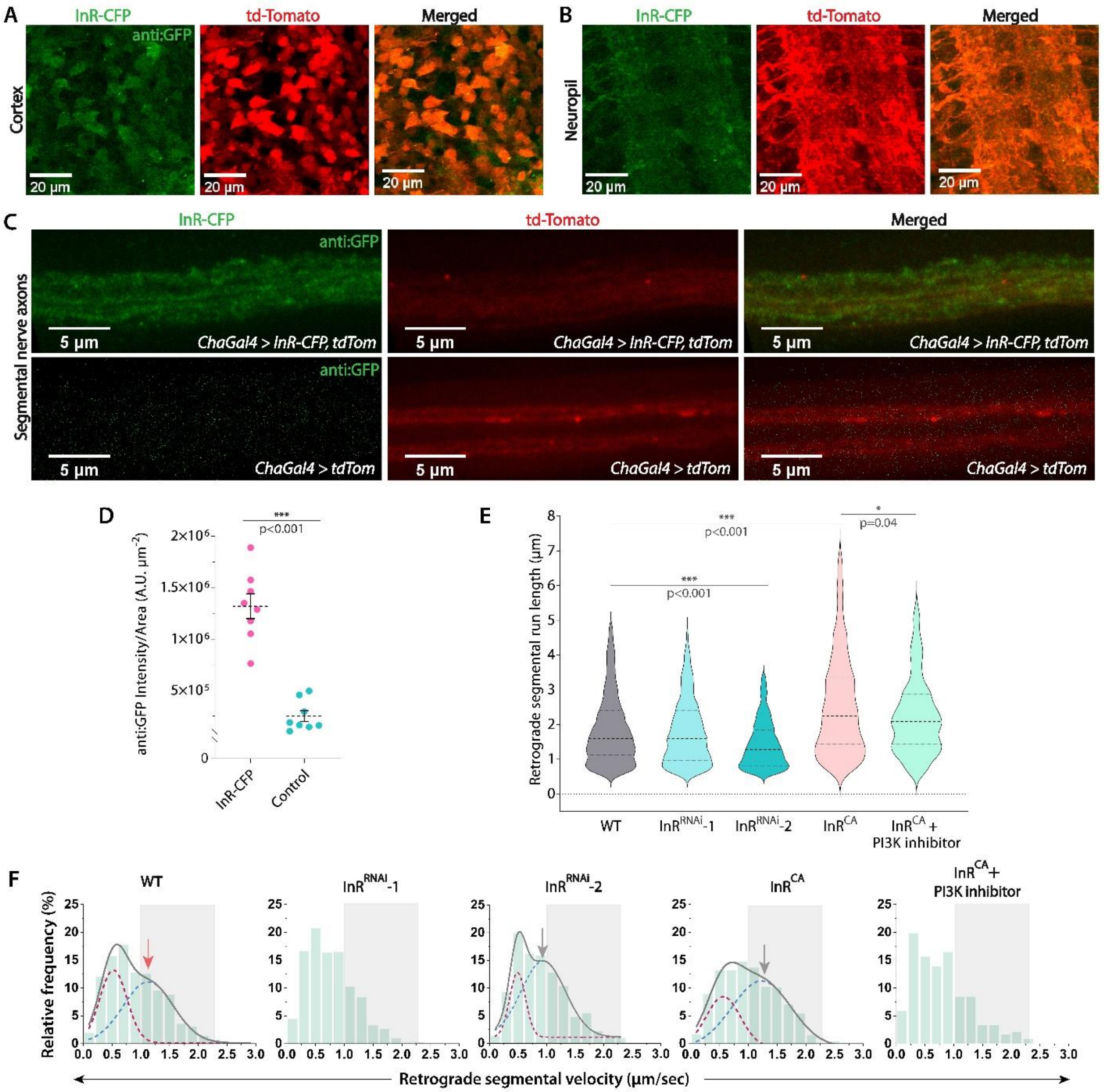


**Supplementary Figure 3: InR-mediated signalling contributes to the axonal transport of Rab4 vesicles.** (A-C) Representative images of InR staining (green) using anti-GFP antibody in the cortex (A), synaptic (B) region, and segmental nerves (C) of the ventral nerve chord expressing InR-CFP and tdTomato in cholinergic neurons of a third instar Drosophila larvae. (D)

Quantification of InR staining (anti-GFP) in the segmental nerves expressing both InR-CFP and tdTomato compared to the segmental nerves expressing only tdTomato. (E-F) Retrograde segmental run length (E) and retrograde segmental velocity (F) of Rab4 vesicles in the wild-type control (WT) and different InRRNAi and InRCA (constitutively active mutant of insulin receptor) overexpression backgrounds at 90 hours AEL (n>200 runs, N = 3-5 larvae each). The cumulative distribution (grey) as a sum of two Gaussians (maroon and blue dotted lines) highlights slow (maroon) and fast-moving (blue) populations (See methods for details). Grey box marks the fast-moving (≥ 1.5 μm/sec) runs. The pairwise significance of difference was estimated using the Mann-Whitney U-test.

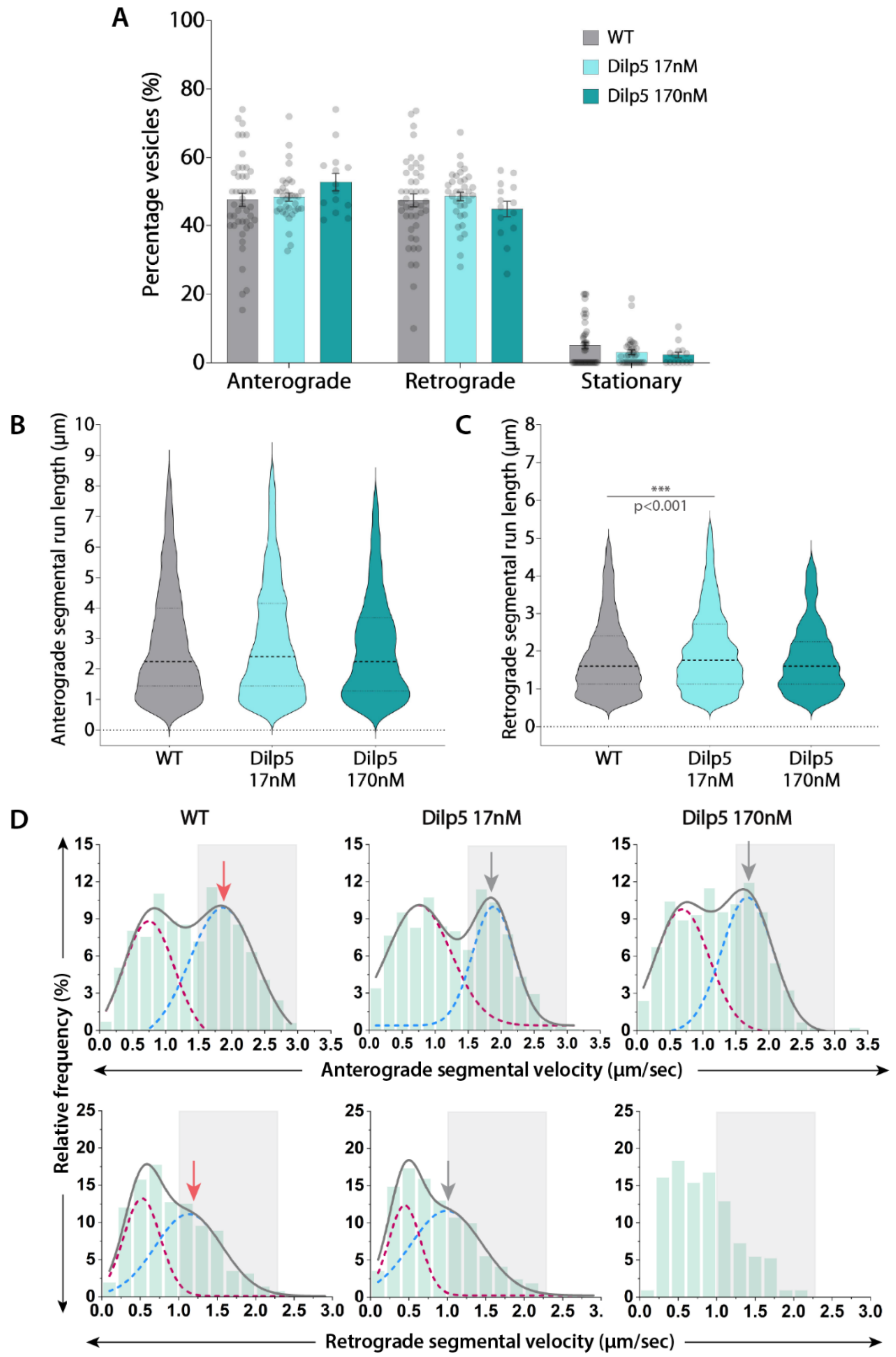


**Supplementary Figure 4: Acute Dilp5 stimulation does not increase the anterograde fraction, velocity, and run length of Rab4 vesicles in the axons.** (A) Relative distribution of the Rab4 vesicle movement (n≥14 segmental nerves, N = 3-5 larvae) in the wild-type control without insulin (WT) and in the presence of different concentrations of Drosophila insulin-like peptide (Dilp5) at 90 hours AEL. The pairwise significance of difference was estimated using the Mann-Whitney U-test. (B, C) Anterograde (B) and retrograde (C) segmental run length (μm) of Rab4 vesicles in wild-type control (WT) and different treatment backgrounds (n>800 runs, N=3-5 larvae each). The pairwise significance of difference was estimated using the Mann-Whitney U-test . (D) Anterograde (top row) and retrograde (bottom row) segmental velocity distributions of Rab4 vesicles in wild-type control (WT) and different treatment backgrounds (n>800 runs; N=3-5 larvae). The cumulative distribution (grey) as a sum of two Gaussians (maroon and blue dotted lines) to highlight slow (maroon) and fast-moving (blue) populations (See methods for details). Grey box marks the fast-moving runs (≥1.5 µm/sec for anterograde and ≥1.0 µm/sec for retrograde).

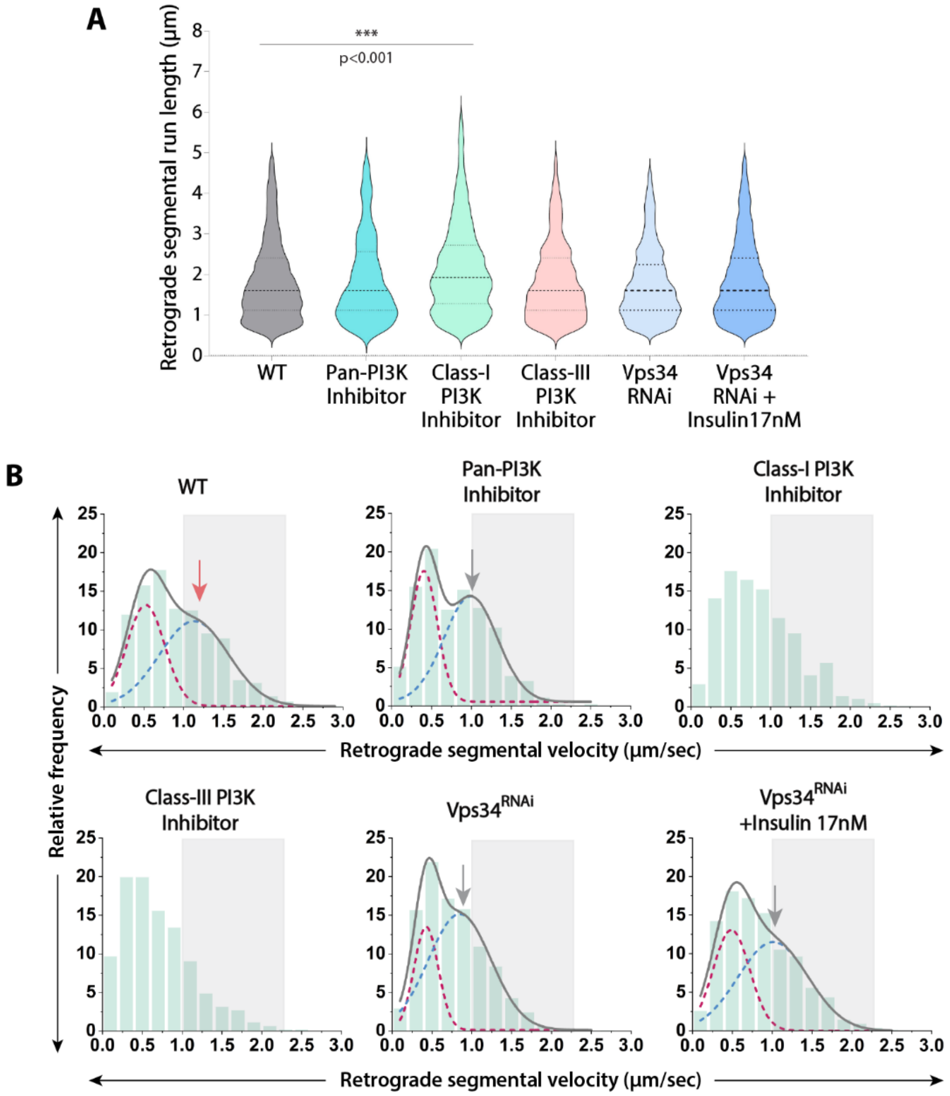


**Supplementary Figure 5: Effects of acute inhibition and knockdown of Class-III PI3K/Vps34 on retrograde movement parameters of Rab4 vesicles in the axon.** (A, B) Retrograde segmental run length (A) and retrograde segmental velocity distributions (B) of Rab4 vesicles in the wild-type control (WT), in the presence of different class-specific PI3K inhibitors, and Vps34RNAi backgrounds in the absence and presence of insulin at 90 hours AEL (n>300 runs, N=3-5 larvae each). The cumulative distribution (grey) as a sum of two Gaussians (maroon and blue dotted lines) highlights slow (maroon) and fast-moving (blue) populations (See methods for details). Grey box marks the fast-moving (≥ 1.0 µm/sec) runs. The pairwise significance of difference was estimated using the Mann-Whitney U-test.

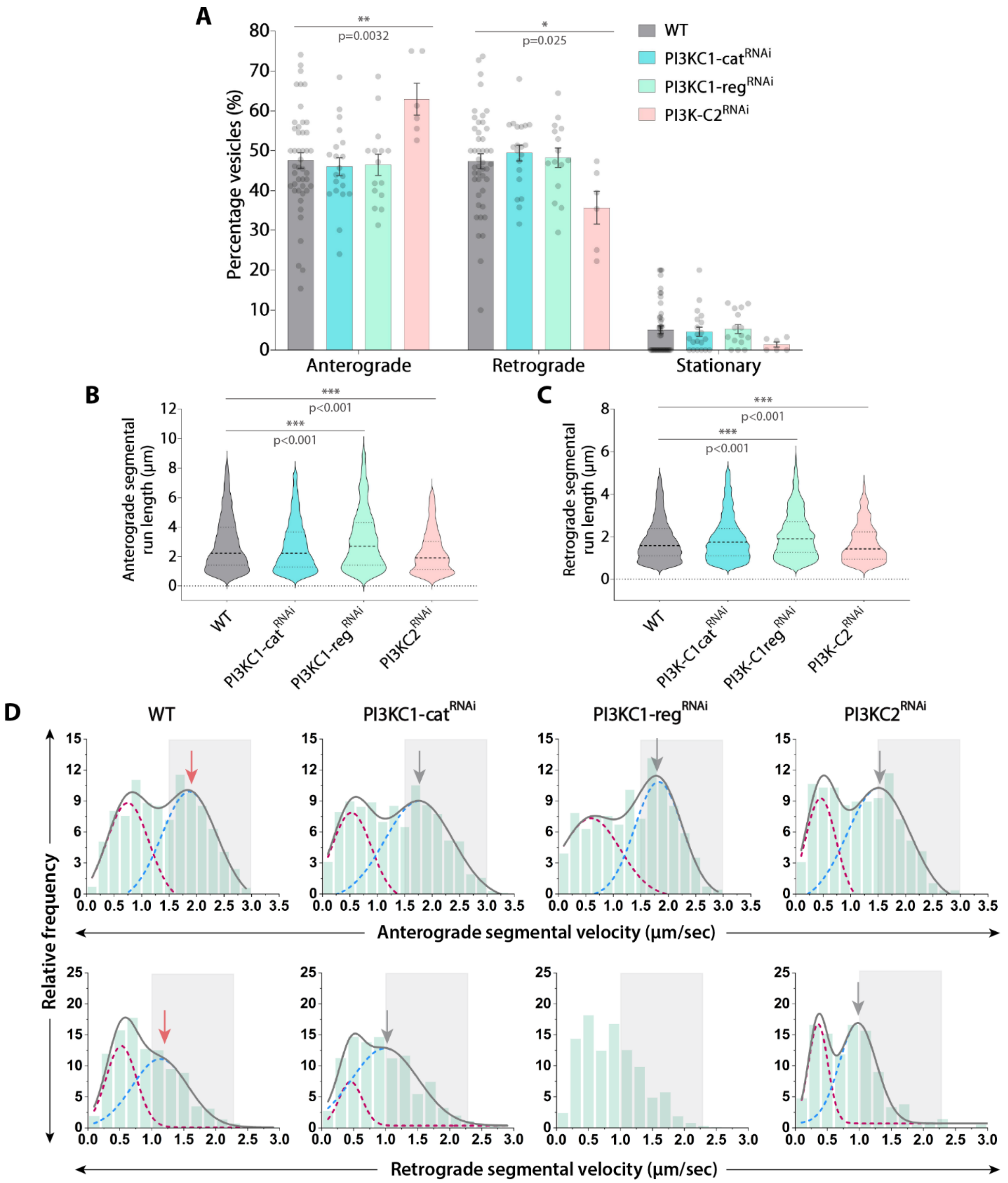


**Supplementary Figure 6: Cell-specific knockdown of Class-I and Class-II PI3Ks does not reduce anterograde fraction of Rab4 vesicles in the axons.** (A) Relative distribution of the Rab4 vesicle movement in the wild-type control (WT) and different class-specific PI3K$^{RNAi}$ backgrounds at 90 hours AEL (n>6 segmental nerves, N=3-5 larvae each). The pairwise significance of difference was estimated using the Mann-Whitney U-test. (B, C) Anterograde (B) and retrograde (C) segmental run length (µm) of Rab4 vesicles in the wild-type control (WT) and different class-specific $PI3K^{RNAi}$ backgrounds (n>200 runs, N=3-5 larvae each). The pairwise significance of difference was estimated using the Mann-Whitney U-test. (D) Anterograde (top row) and retrograde (bottom row) segmental velocity distributions of Rab4 vesicles in the wild-type control (WT) and class-specific $PI3K^{RNAi}$ knockdown backgrounds (n>200 runs, N=3-5 larvae each).The cumulative distribution (grey) as a sum of two Gaussians (maroon and blue dotted lines) to highlight slow (maroon) and fast-moving (blue) populations (See methods for details). Grey box marks the fast-moving runs (≥1.5 μm/sec for anterograde and ≥1.0 μm/sec for retrograde).

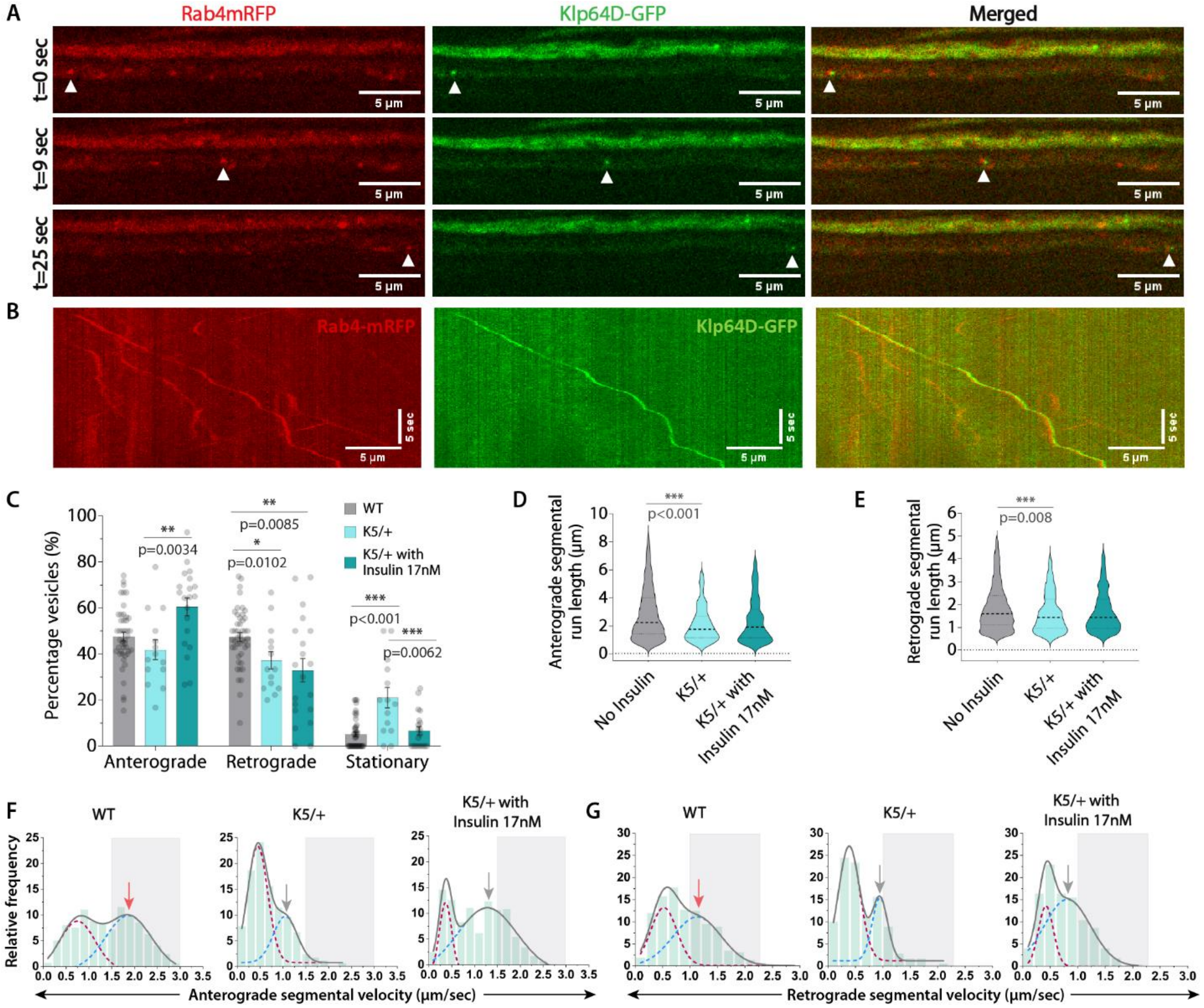


**Supplementary Figure 7: Kinesin-2 does not regulate the increase in anterograde fraction of Rab4 vesicles in the axons downstream of insulin signaling.** (A, B) Simultaneously acquired dual-channel time-lapse images of segmental nerve (A) and kymographs (B) depict comigration of genetically-encoded Rab4mRFP (red) and Klp64DGFP (green, KIF3A ortholog) in cholinergic axons. (C) Relative distribution of the Rab4 vesicle movement in the wild-type control (WT) and kinesin-2 mutant, K5, in the heterozygous background in the absence and presence of insulin at 90 hours AEL (n≥14 segmental nerves, N=3-5 larvae each). The pairwise significance of difference was estimated using the Mann-Whitney U-test. (D, E) Anterograde (D) and retrograde (E)

segmental run length (µm) of Rab4 vesicles in the wild-type control (WT) and different genetic backgrounds (n>150 runs, N=3-5 larvae each). The pairwise significance of difference was estimated using the Mann-Whitney U-test. (F, G) Anterograde (F) and retrograde (G) segmental velocity distributions of Rab4 vesicles in the wild-type control (WT) and different genetic backgrounds (n>150 runs, N=3-5 larvae each) at 90 hours AEL. The cumulative distribution (grey) as a sum of two Gaussians (maroon and blue dotted lines) to highlight slow (maroon) and fast-moving (blue) populations (See methods for details). Grey box marks the fast-moving runs (≥1.5 µm/sec for anterograde and ≥1.0 µm/sec for retrograde).

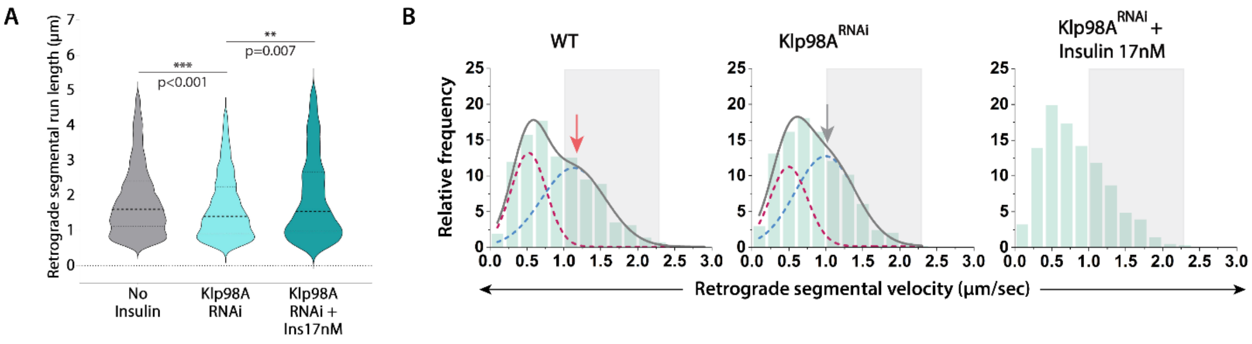


**Supplementary Figure 8: Effect of Klp98A knockdown on the retrograde movement parameters of Rab4 vesicles in the axons.** (A, B) Retrograde segmental run length (A) and retrograde segmental velocity distributions (B) of Rab4 vesicles in the wild-type control (WT) and Klp98A$^{RNAi}$ in the absence and presence of insulin (n>400 runs, N=4-6 larvae each) at 90 hours AEL. The cumulative distribution (grey) as a sum of two Gaussians (maroon and blue dotted lines) highlights slow (maroon) and fast-moving (blue) populations (See methods for details). Grey box marks the fast-moving (≥ 1.0 µm/sec) runs. The pairwise significance of difference was estimated using the Mann-Whitney U-test.

**Table S1.**

Percentage of fast- and slow-moving vesicles in anterograde and retrograde directions across conditions, pharmacological treatments, and genetic perturbations.

| Condition/Treatment/Perturbation | Anterograde % | | Retrograde % | |
|---|---|---|---|---|
| | 0-1.6µm/sec | >1.6µm/sec | 0-1µm/sec | >1µm/sec |
| **72 hours AEL (After Egg Laying)** | 76.92 | 23.08 | 73.46 | 26.54 |
| **80 hours AEL** | 59.84 | 40.16 | 68.59 | 31.41 |
| **No insulin/90 hours AEL** | 56.82 | 43.18 | 60.05 | 39.95 |
| **Insulin 1.7nM** | 48.33 | 51.67 | 50.53 | 49.47 |
| **Dilp2 17nM** | 51.34 | 48.66 | 59.98 | 40.02 |
| **Dilp2 170nM** | 41.46 | 58.54 | 47.84 | 52.16 |
| **Dilp5 17nM** | 62.32 | 37.68 | 64.63 | 35.37 |
| **Dilp5 170nM** | 68.83 | 31.17 | 67.42 | 32.58 |
| **LY294002 50uM** | 78.61 | 21.39 | 68.57 | 31.43 |
| **HS173 50nM** | 64.07 | 35.93 | 66.15 | 33.85 |
| **SAR405 50uM** | 81.44 | 18.56 | 78.38 | 21.62 |
| **InR$^{CA}$** | 42.65 | 57.35 | 51.54 | 48.46 |
| **InR$^{CA}$ + LY294002 560uM** | 70.2 | 29.8 | 71.73 | 28.27 |
| **InR$^{RNAi}$-1** | 72.94 | 27.06 | 74.4 | 25.6 |
| **InR$^{RNAi}$-2** | 72.56 | 27.44 | 62.66 | 37.34 |
| **Class-I PI3K cat$^{RNAi}$** | 59.16 | 40.84 | 57.08 | 42.92 |
| **Class-I PI3K reg$^{RNAi}$** | 57.13 | 42.87 | 64.04 | 35.96 |
| **Class-II PI3K$^{RNAi}$** | 70.11 | 29.89 | 64.93 | 35.07 |
| **Vps34$^{RNAi}$** | 67.52 | 32.48 | 73.31 | 26.69 |
| **Vps34$^{RNAi}$ + Insulin 17nM** | 63.66 | 36.34 | 67.29 | 32.71 |
| **Klp64D$^{K5}$** | 96.38 | 3.62 | 84.44 | 15.56 |
| **Klp64D$^{K5}$ + Insulin 17nM** | 77.8 | 22.2 | 75.07 | 24.93 |
| **Klp98a$^{RNAi}$** | 84.44 | 15.56 | 66.41 | 33.59 |
| **Klp98a$^{RNAi}$ + Insulin 17nM** | 73.72 | 26.28 | 68.55 | 31.45 |

**Supplementary Video S1.**

Axonal transport of Rab4mRFP vesicles in the distal axons of *Drosophila* cholinergic neurons at 72, 80, and 90 hours AEL. Widefield time-lapse imaging was performed at 10fps, and the movies are played at 50fps. Duration of all the representative movies is 30sec.

**Supplementary Video S2.**

Effect of various genetic and pharmacological perturbations of insulin signalling on the axonal transport of Rab4mRFP vesicles in the distal axons of *Drosophila* cholinergic neurons at 90 hours AEL. Widefield time-lapse imaging was performed at 10fps, and the movies are played at 50fps. Duration of all the representative movies is 30sec.

**Supplementary Video S3.**

Effect of acute stimulation of human insulin and Dilp2 on the axonal transport of Rab4mRFP vesicles in the distal axons of *Drosophila* cholinergic neurons at 90 hours AEL. Widefield time-lapse imaging was performed at 10fps, and the movies are played at 50fps. Duration of all the representative movies is 30sec.

**Supplementary Video S4.**

Effect of acute inhibition of PI3Kinases on the axonal transport of Rab4mRFP vesicles in the distal axons of *Drosophila* cholinergic neurons at 90 hours AEL. Widefield time-lapse imaging was performed at 10fps, and the movies are played at 50fps. Duration of all the representative movies is 30sec.

**Supplementary Video S5.**

Effect of cell-specific knockdown of Class-III PI3K/Vps34 with and without acute insulin stimulation on the axonal transport of Rab4mRFP vesicles in the distal axons of *Drosophila* cholinergic neurons at 90 hours AEL. Widefield time-lapse imaging was performed at 10fps, and the movies are played at 50fps. Duration of all the representative movies is 30sec.

**Supplementary Video S6.**

Simultaneous dual-color time-lapse imaging performed using Spinning-disc confocal microscopy shows comigration of genetically-encoded PI(3)P biosensor (2xFYVE-GFP) and Rab4mRFP in the distal axons of *Drosophila* cholinergic neurons. Data was collected at 8-9fps, and movies are played at 50fps. Duration of the representative movie is 20sec.

**Supplementary Video S7.**

Simultaneous dual-color time-lapse imaging performed using Spinning-disc confocal microscopy shows comigration of 2xFYVE-GFP and Rab4mRFP in the distal axons of *Drosophila* cholinergic neurons. Particle 1 (green) and Particle 2 (cyan) are representative examples of particles with a low and high 2xFYVE-GFP/Rab4mRFP intensity ratio (as plotted in Figure 6F), respectively. Data was collected at 8-9fps, and movies are played at 50fps. Duration of the representative movie is 20sec.

**Supplementary Video S8.**

Simultaneous dual-color time-lapse imaging performed using Spinning-disc confocal microscopy shows comigration of Klp98a-GFP and Rab4mRFP in the distal axons of *Drosophila* cholinergic neurons. Data was collected at 8-9fps, and movies are played at 50fps. Duration of the representative movie is ~20sec.

**Source Data S1. (separate file)**

Raw data values for volume and intensity of Brp per hemisegment and segment-wise in A3-A6 from 72-90 hours AEL (Fig.1 D-E and fig. S1B-C) is organized in different sheets in the spreadsheet.

**Source Data S2. (separate file)**

Raw data values for volume and intensity of soluble GFP marked by *ChaGal4* per hemisegment in A3-A6 (Fig.1 G-H) is organized in different sheets in the spreadsheet.

**Source Data S3. (separate file)**

Raw data values for density (intensity per unit volume) of Rab4 and Bruchpilot per hemisegment in A3-A6 (Fig. 2B).

**Source Data S4. (separate file)**

Raw data values of percentage of Rab4 vesicles in the anterograde, retrograde, and stationary categories (fraction) at different developmental timepoints, in different conditions, or after a pharmacological or genetic perturbation (Fig. 2D, 3A, 4A, 5A, 5D, 7C and fig. S4A, S6A, S7C).

**Source Data S5. (separate file)**

Raw data values of segmental run length (µm) of Rab4 vesicles in the anterograde and retrograde direction at different developmental timepoints, in different conditions, or after a pharmacological or genetic perturbation (Fig. 2E, S2A, 3B, 4B-C, 5B, 5E, 7D and fig. S3E, S4B-C, S5A, S6B-C, S7D-E, S8A).

**Source Data S6. (separate file)**

Raw data values of segmental velocity (µm/sec) of Rab4 vesicles in the anterograde and retrograde direction at different developmental timepoints, in different conditions, or after a pharmacological or genetic perturbation (Fig. 2F, S2B, 3C, S3F, 4D, S4D, 5C, 5F, S5B, S6D, 7E, S7F, and S8B).

**Source Data S7. (separate file)**

Raw data values of anti-GFP intensity per unit area values (A.U. $\mu m^{-2}$) in the segmental nerves axons for control and InR-CFP overexpression background (fig. S3C-D).

**Source Data S8. (separate file)**

Raw data values of percentage colocalized vesicles of the total (Rab4mRFPand 2xFYVE-GFP) in different pharmacological perturbations in wandering third instar larvae (Fig. 6C-D) and at different developmental timepoints (Fig. 8C-D).

**Source Data S9. (separate file)**

Raw data values of Rab4mRFP and 2xFYVE-GFP intensities (A.U.) on single vesicles and their average velocities (µm/sec) in the distal axons of cholinergic neurons (Fig. 6E-F).

**Source Data S10. (separate file)**

Raw data values of percentage colocalized vesicles of the total (Rab4mRFP and Klp98A-GFP) in different pharmacological perturbations in wandering third instar larvae (Fig. 7F-G) and at different developmental timepoints (Fig. 8A-B).
